# Protease activities of vaginal *Porphyromonas* species disrupt coagulation and extracellular matrix in the cervicovaginal niche

**DOI:** 10.1101/2021.07.07.447795

**Authors:** Karen V. Lithgow, Vienna C.H. Buchholz, Emily Ku, Shaelen Konschuh, Ana D’Aubeterre, Laura K. Sycuro

## Abstract

*Porphyromonas asaccahrolytica* and *Porphyromonas uenonis* are frequently isolated from the human vagina and are linked to bacterial vaginosis and preterm labour. However, little is known about the pathogenesis mechanisms of these bacteria. The related oral opportunistic pathogen, *Porphyromonas gingivalis,* is comparatively well-studied and known to secrete numerous extracellular matrix-targeting proteases. Among these are the gingipain family of cysteine proteases that drive periodontal disease progression and hematogenic transmission to the placenta. Given their phylogenetic relatedness, we hypothesized that vaginal *Porphyromonas* species possess gingipain-like protease activity targeting host extracellular matrix in the female reproductive tract. In this study, we demonstrate that vaginal *Porphyromonas* species degrade type I collagen (cervix), type IV collagen (chorioamnion/placenta), and fibrinogen, but not through the activity of gingipain orthologs. Bioinformatic queries identified 5 candidate collagenases in each species, including serine, cysteine and metalloproteases, with signal peptides directing them to the extracellular environment. Inhibition assays revealed both species secrete metalloproteases that degrade collagen and casein, while *P. asaccharolytica* also secretes a metalloprotease that degrades fibrinogen. Phylogenetic analysis of the predicted collagen-degrading metalloprotease revealed an orthologous relationship with the *P. gingivalis* endopeptidase PepO. Cloning and expression of *P. asaccharolytica* PepO confirmed this protein’s collagenase and caseinase activities, which have not previously been attributed to PepO homologs in other bacteria. Altogether, this description of the first known virulence factor in *Porphyromonas* species colonizing the human vagina sheds light on their potential to alter the structural integrity and homeostasis of reproductive tissues.

**Importance:** *Porphyromonas* species are common inhabitants of the vaginal microbiome, but their presence has been liked to adverse health outcomes for women, including bacterial vaginosis and preterm birth. We determined that *P. asaccharolytica* and *P. uenonis* secrete broad-acting proteases capable of freely diffusing within the cervicovaginal niche and degrading important components of host tissues, namely the extracellular matrix. We show that secreted *Porphyromonas* proteases degrade collagens that are enriched within the cervix (type I) and chorioamniotic membranes (type IV). Furthermore, these *Porphyromonas* proteases can also degrade fibrinogen and inhibit clot formation. These activities can be partially attributed to a metalloprotease that exhibits broad-acting protease activity and is distantly related to the *P. gingivalis* endopeptidase PepO. This initial characterization of virulence activities in vaginal *Porphyromonas* species highlights their potential to harm human pregnancy through clotting disruption, fetal membrane weakening, and premature cervical remodeling.

## Introduction

The vaginal microbiome of healthy reproductive-age women is typically characterized by low species diversity, with *Lactobacillus* dominating the vaginal and ectocervical niches of the lower genital tract (1, 2). A community shift towards high species diversity, with overgrowth of anaerobic bacteria, is associated with increased risk of bacterial vaginosis (BV) (2, 3), acquisition and transmission of sexually transmitted infections (4–6), preterm birth (7–9), and cervical cancer (10–12). Intriguingly, some women harbouring a diverse cervicovaginal microbiome are healthy and asymptomatic (2), suggesting a need to untangle how specific species contribute to poor outcomes. Among BV-associated bacteria, Gram-negative anaerobic rods corresponding to *Prevotella* and black-pigmented *Porphyromonas* species are frequently detected in vaginal samples and significantly associated with the *Bacteroides* morphotype from Nugent scoring (13). While *Prevotella* is an abundant, species-rich and relatively well- studied vaginal clade, comparatively little is known about *Porphyromonas* species inhabiting the human vagina. No *Porphyromonas* species is currently thought to be specific to the human urogenital tract, but *P. asaccharolytica*, *P. uenonis, P. bennonis* and *P. somerae* (in decreasing order of cervicovaginal microbiome citation frequency) exhibit a preference for these niches (3, 14, 15). *P. asaccharolytica* and *P. uenonis* colonize the vagina in 15–50% of healthy women and although their prevalence and abundance increases with BV, they are typically considered low abundance taxa (13, 16–20). Recent studies show these species are predictors of spontaneous preterm labour (9, 21), pelvic inflammatory disease (22, 23), human papillomavirus (HPV) infections progressing to cervical neoplasia (24, 25), and uterine cancer (26, 27). Thus, an improved understanding of the functional capacity of vaginal *Porphyromonas* species is needed.

To date, the only *Porphyromonas* species that has been well-characterized is *P. gingivalis*, a low abundance species in the oral microbiome of both healthy patients and those with gingivitis (28, 29). As a keystone species driving oral (plaque) biofilm formation and periodontal disease progression (30, 31), *P. gingivalis* contributes to local tissue destruction directly and indirectly through the induction of inflammatory processes (32, 33). *P. gingivalis* can also disseminate via the bloodstream to distal infection sites such as the endocardium and joints (34, 35). During pregnancy, *P. gingivalis* has been isolated from the placenta and amniotic fluid of women who delivered preterm (36–38), and in mouse infection models, *P. gingivalis* induces preterm labour via inflammatory activation of the chorioamniotic membranes (34, 39, 40). Pathogenesis mechanisms contributing to these outcomes include a wide array of proteolytic activities carried out by numerous secreted proteases. Among these are the gingipain family of cysteine proteases that drive periodontal disease progression (41), hematogenic transmission to the placenta (29, 40, 42, 43) and preterm labour induction in mice (40). The gingipains degrade many extracellular matrix components, including collagen (44, 45), and amplify their effects by activating and upregulating host matrix metalloproteases (MMPs) that also degrade collagen (46, 47). Furthermore, gingipains can degrade immune factors including immunoglobulins (44, 48), complement components (49, 50), cytokines (51, 52), clotting factors (53, 54), and antimicrobial peptides (55), giving rise to a favourable immune environment for *P. gingivalis* colonization.

Proteolytic activity has been previously detected in vaginal fluid from patients with BV (56–58) and characterized in clinical isolates of BV-associated bacteria (59, 60). In fact, collagenase (gelatinase) and caseinase activity of *P. asaccharolytica* (formerly *Bacteroides asaccharolyticus*) was previously reported in screens of *Bacteroides* species detected in human infections (61) and clinical isolates from reproductive tract infections (60). However, further characterization of the enzymes responsible was not conducted, and proteolytic activity of *P. uenonis* has yet to be explored. Given their phylogenetic relatedness and epidemiological similarity, exhibiting high prevalence, low abundance and association with disease, we sought to determine whether vaginal *Porphyromonas* species possess the broad-acting proteolytic virulence activity of *P. gingivalis*. In this study we show that *P. asaccharolytica* and *P. uenonis* are both capable of degrading several extracellular matrix components found within the female genital tract. Our study furthermore reveals differences between the species, suggesting preterm birth-associated *P. asaccharolytica* may secrete more enzymes that contribute collagenase activity. Finally, we report the first virulence factor identified and functionally characterized in *P. asaccharolytica* – a metalloprotease that is highly conserved in *P. uenonis* and more distantly related to to the PepO endopeptidase in *P. gingivalis* and other *Porphyromonas* species (62, 63). We demonstrate this protein exhibits broad-acting proteolytic capacity, which may make it a key microbial virulence factor in the pathogenesis of reproductive health conditions and gynecological cancers.

## Materials & Methods

### Bacterial Strains and Growth Conditions

*Porphyromonas asaccharolytica* CCUG 7834 (type strain, identical to DSM 20707, ATCC 25260 and JCM 6326), *Porphyromonas uenonis* CCUG 48615 (type strain, identical to DSM 23387, ATCC BAA-906 and JCM 13868), *Porphyromonas gingivalis* W50 ATCC 53978 and *Lactobacillus crispatus* CCUG 42897 were cultured anaerobically on 1.5% brucella agar (BD Biosciences, Franklin Lakes, MD) supplemented with 5% defibrinated sheep’s blood (Dalynn Biologicals, Calgary, AB). For liquid cultivation, supplemented brain heart infusion (sBHI) was prepared by supplementing BHI (BD) with 2% gelatin (BD), 1% yeast extract (ThermoFisher Scientific, Burnaby, BC), 0.8% dextrose (BD) and 0.1% starch (ThermoFisher). Solid and liquid cultivation was conducted at 37°C in an AS-580 anaerobic chamber (Anaerobe Systems, Morgan Hill, CA). Bacterial suspensions were prepared by harvesting cells from solid medium after growth for 16-24 hours (*P. gingivalis, L. crispatus*) or 36-48 hours (*P. asaccharolytica*, *P. uenonis)* and resuspending cells in sBHI, sBHI (no gelatin), BHI (no supplements) or PBS. Optical density at 600 nm (OD600) of bacterial suspensions was measured with a Genesys 300 visible spectrophotometer (ThermoFisher Burnaby, BC) and colony forming units per mL (cfu/mL) was calculated using empirically determined cfu/mL/OD600nm for each strain. Serial dilution spot plating was used to verify the cfu/mL of the starting suspensions. Cell-free supernatants (SNs) were harvested during late-log to early stationary phase for each species (*P. asaccharolytica* OD600 1.3-1.8; *P. uenonis* OD600 0.8-1.2). Liquid cultures were centrifuged at 10,000 × g for 10 minutes at room temperature before the supernatant was filter sterilized (0.2 µm; Pall Laboratory, Mississauga, ON) and stored at -20°C.

### Collagenase Assays

Cell suspensions or cell-free supernatants were tested for collagenase activity with the EnzChek Gelatinase/Collagenase assay kit (Invitrogen, Carlsbad, CA) using fluorescein- labelled DQ^TM^ gelatin conjugate (type I collagen, Invitrogen) or a type IV DQ^TM^ collagen conjugate from human placenta (Invitrogen). Reactions were prepared in technical triplicate or quadruplicate by mixing 20 µL of substrate at 0.25 mg/mL with 80 µL of reaction buffer and 100 µL of bacterial suspension, cell- free supernatant or media in black optical bottom 96-well plates (Greiner Bio-One, Monroe, NC). Using a Synergy H1 microplate reader (BioTek, Winooski, VT), plates were incubated at 37°C in atmospheric conditions. Kinetic fluorescence reads were measured at 485 nm excitation/527 nm emission every three minutes over two hours, or every thirty minutes over eighteen hours, for cell suspension and cell- free supernatant assays, respectively. Prior to fluorescence reads, plates were shaken for seven seconds. The mean fluorescence readings of the negative control (substrate in sBHI media) were subtracted from experimental wells and relative fluorescence units (RFU) was plotted over time, with negative values adjusted to zero.

### Zymography

The total protein content of *Porphyromonas* cell-free supernatants was determined using a bicinchoninic acid microplate assay (BCA; Pierce, Rockford, IL). Cell-free supernatants were diluted to 8 mg/mL and 5 µL of sample was combined with 5 µL of Novex^TM^ Tris-Glycine SDS Sample Buffer (Invitrogen) to load 40 µg per well. Samples were separated on Novex^TM^ 10% Zymogram Plus (Gelatin; Invitrogen) protein gels at a constant voltage of 125 V in Novex^TM^ 1X Tris-Glycine Running Buffer (Invitrogen). After separation, gels were incubated in Novex^TM^ 1X Renaturing Buffer for 30 minutes at room temperature with gentle agitation, followed by two consecutive incubations in Novex^TM^ 1X Developing Buffer: room temperature for 30 minutes and 37°C for 16 hours. Gels were then stained in Coomassie brilliant blue R-250 solution (Fisher Scientific) and de-stained in 5% (vol/vol) methanol/7.5% (vol/vol) acetic acid in distilled water.

### Caseinase Assay

*Porphyromonas* cell-free supernatants were tested for general protease activity using fluorescein-labelled casein (FITC-Casein, ThermoFisher) as a substrate. Reactions were prepared in technical triplicate by mixing 20 µL of substrate at 50 µg/mL with 80 µL of Tris-buffered saline and 100 µL of cell-free supernatant in black optical bottom 96-well plates (Greiner Bio-One). Fluorescence plate reader measurements were performed as described above by measuring 485 nm excitation/527 nm emission every ten minutes over five hours.

### Casein Plate Assay

Casein plates were prepared by autoclaving three solutions: 30 g/L instant skim milk power (Pacific Dairy), 19 g/L brain heart infusion (BHI, BD Biosciences) and 30 g/L agar (Fisher Scientific). The solutions were combined in equal volume and 10 mL was added to 100×15 mm petri dishes (Fisher Scientific) to solidify at room temperature. Bacterial suspensions were prepared in PBS and 5 µL was spotted onto casein agar plates. Zones of clearance were measured for each spot after incubating the plates at 37°C under anaerobic conditions for three to six days.

### Clotting Assays

Bacterial strains were harvested from solid medium and suspended in 13 mM sodium citrate. Suspensions were centrifuged at 10,000 × g for 7 minutes at room temperature and resuspended in 13 mM sodium citrate. Duplicate cell suspensions, or cell-free controls, were incubated with 50 µL of sterile filtered bovine plasma (Quad Five, Ryegate, MT) for 30 minutes at 37°C under anaerobic conditions. After incubation, samples were blinded and centrifuged at 10,000 × g for seven minutes at room temperature, and 50 µL of HEMOCLOT thrombin time reagent (Hyphen Biomed, Neuville-sur-Oise, France) was added to each reaction. The clotting time for each sample was estimated using a stereo microscope to directly visualize clot formation, indicated by the presence of white precipitate, tendril formation or increased viscosity of the samples. The clotting time for samples that did not form visible clots was recorded as 1800 seconds. To further evaluate final clot size, samples were transferred to a clear 96-well plate and OD (405 nm) was measured for the entire well using the well area scan feature of the Synergy H1 microplate reader (BioTek). The average OD405 nm for each well was blanked against duplicate thrombin-free controls for each experimental sample type.

### Fibrinogen Degradation Assay

*Porphyromonas* cell suspensions (10^7^ cfu/reaction), sBHI media controls or *Porphyromonas* cell-free supernatants (240 µg of total protein) were incubated with 120 µg of human fibrinogen (Sigma-Aldrich, St. Louis, MO). Reaction mixtures were incubated at 37°C under anaerobic conditions or in an atmosphere of 5% CO2 for cell suspension or cell-free supernatants, respectively. Samples from each time point (cell suspensions: 0, 2, 18, 24 hours; supernatants: 0, 2, 24, 48 hours) were collected, mixed 1:1 with Novex^TM^ 2X sample buffer (Invitrogen) with dithiothreitol (DTT, Fisher Scientific), heated at 95°C for 10 minutes and separated on Novex^TM^ 10% Tris-Glycine polyacrylamide pre-cast gels (Invitrogen) at a constant voltage of 180 V. Gels were stained in Coomassie brilliant blue R-250 (0.25% w/v) solution (Fisher Scientific) and de-stained in 5% (vol/vol) methanol/7.5% (vol/vol) acetic acid in H2O. Fibrinogen degradation was evaluated qualitatively by visualization of fibrinogen α chain (63.5 kDa), β chain (56 kDa) and γ chain (47 kDa) between experimental and control samples over the time-course.

### Protease Inhibition Assays

Protease inhibitors were incorporated into collagenase, caseinase and fibrinogen degradation assays, and working solutions were prepared in reaction buffer or TBS. The metalloprotease inhibitor, 1,10-Phenanthroline (Invitrogen), was prepared as a 2 M stock solution in ethanol and diluted to working concentrations of 0.2, 0.02 or 0.002 mM. Iodoacetamide (G-Biosciences, St. Louis, MO, or Sigma-Aldrich) was prepared as a 10 mM stock solution in HyPure H2O (Cytiva Life Sciences, Marlborough, MA) and used at working concentrations of 0.4, 0.04 or 0.004 mM to inhibit cysteine protease activity. The serine protease inhibitor, aprotinin (Roche, Mississauga, ON) was prepared as a 0.1 mM stock solution in HyPure H2O (Cytiva Life Sciences) and used at working concentrations of 0.01, 0.001 or 0.0001 mM.

### Bioinformatic Analyses

*Gingipain Ortholog Queries.* Gingipain amino acid sequences (RgpA (PGN_1970; P28784), RgpB (PGN_1466; P95493) and Kgp (PGN_1728; B2RLK2) were obtained from *Porphyromonas gingivalis* ATCC 33277 through UniProtKB. Each sequence was queried against all available *P. asaccharolytica* (DSM 20707; PR426713P-I) and *P. uenonis* (DSM 23387; 60-3) genomes using the ‘Selected Genomes’ protein Basic Local Alignment Search Tool (BLAST; default settings) in the IMG/MER database to identify potential orthologs (Supplemental Figure 1A). Each gingipain AA sequence was also submitted to the Pfam database and all Pfam IDs and names were recorded. Gingipain Pfam IDs were searched against all available *P. asaccharolytica* (DSM 20707; PR426713P-I) and *P. uenonis* (DSM 23387 [IMG Genome ID 2585427891 and 2528311143]; 60-3) strains using the advanced gene search function in the IMG/MER database (Supplemental Figure 5A). All sequences returned by these IMG/MER BLAST and Pfam ID searches were used to query *P. gingivalis* ATCC 33277 in a reciprocal BLAST search with the National Center for Biotechnology Information (NCBI) protein BLAST tool (Supplemental Figure 1A). Select hits were aligned with the *P. gingivalis* gingipains across their full length using the EMBL-EMI Clustal Omega alignment tool (64).

#### Candidate Collagenase Queries

Peptidases from *P. asaccharolytica* CCUG 7834 and *P. uenonis* 60-3 were identified by searching for entries with MEROPS peptidase annotations in UniProt (65). For the microbial peptidases from *P. asaccharolytica* and *P. uenonis*, the following enzyme information was exported from UniProtKB for each entry: gene ontology (biological process and molecular function), MEROPs, Pfam, PANTHER, PROSITE, SMART, SUPFAM. Next, a microbial collagenase enzyme number (EC3.4.24.3) was identified in BRENDA (66) and searched against the UniProtKB database (67), generating a list of 3417 entries corresponding to predicted and confirmed microbial collagenases. These microbial collagenase identifiers were cross-referenced against the exported peptidase information from *P. asaccharolytica* and *P. uenonis* to generate a short-list of 14–18 candidate collagenases in *P. asaccharolytica* and *P. uenonis* (Supplemental Figure 1B). Short-list candidates were explored in UniProt and InterPro Scan (68) to eliminate any proteins involved in cell wall synthesis or export machinery and identify the most promising candidates. Sequences were evaluated for the presence of secretion signals using SignalP v.5.0 (69) or integrated information in InterPro. Presence of the IDs: TIGR0483, IPR026444, or PF18962 from TIGR Fam, InterPro and Pfam, respectively, within protein C-termini was indicative of secretion via the type IX secretion system. Since the MEROPs peptidase database only contained information for *P. uenonis* 60-3, IMG BLAST searches were used to identify the corresponding candidate collagenase in the experimental strain used in this study, *P. uenonis* CCUG 48615. Sequence identity and InterPro scans were evaluated for the top hit from each BLAST search in *P. uenonis* CCUG 48615 (Supplemental Table 6).

#### Phylogenetic Analysis

16S rRNA gene sequences were collected using the Integrated Microbial Genomes & Microbiomes (IMG/MER) (70) or National Center for Biotechnology Information (NCBI) databases for each *Porphyromonas* species reported in the human urogenital tract by Acuna-Amador in their comprehensive review (14). Uncultured and recently cultured *Porphyromonas* species were also included (71, 72). Nomenclature choices were informed by the ANI tool within IMG/MER and the Genome Taxonomy Database (GTDB, Release 06-RS202, April 27^th^, 2021) (73). Whenever possible, type strain and near full-length sequences were selected. Multiple sequence alignment was performed with SINA aligner (v.1.2.11) (74) using Silva’s Alignment, Classification and Tree Service (ACT) (75). The phylogenetic tree was computed using RAxML v.8.2.9 (76) with the Gamma model for likelihoods (also through ACT). The tree was edited using the interactive Tree of Life (iTOL (77)). The AA identity of PepO orthologues in each *Porphyromonas* species was queried through BLASTP searches in IMG/MER and NCBI using the *P. asaccharolytica* AA sequence as the query. With the exception of *P. uenonis* 60.3, the AA identity reported corresponds with a hit showing >98% query coverage.

### Construct cloning and *in vitro* transcription/translation

The Poras_0079 (PepO; IMG Gene ID 2504823953) DNA fragment encoding amino acid residues C21 to W682 was PCR amplified from a *P. asaccharolytica* DSM 20707 genomic DNA extraction (Qiagen DNeasy Blood & Tissue Kit, Germantown, MD) using the forward (5’- ATATCCATGGCTTGTAACAAGAAGCAGGAGAATC-3’) and reverse primers (5’- ATATCCCGGGCCAGACCACGACACGCTC-3’). The amplicon was cloned into the pTXTL-T7p14-aH plasmid (replacing alpha hemolysin, Daicel Arbor Biosciences, Ann-Arbor, MI) using NcoI-HF and SmaI (New England Biolabs, Ipswitch, MA). The new plasmid construct (pSLP15) was transformed into *Escherichia coli* DH5α chemically competent cells and prepped using the QIAprep spin miniprep kit (Qiagen). Plasmid concentrations were determined using the Qubit dsDNA broad range assay kit with a Qubit 3 fluorometer (Invitrogen). *In vitro* myTXTL reactions were prepared by combining 5 nM of pSLP15, 1 nM of pTXTL-P70a-T7rnap (expressing the T7 RNA polymerase, Daicel Arbor BioSciences, Ann-Arbor, MI) and 9 µl of myTXTL Sigma 70 Master Mix (Arbor Biosciences) to a final volume of 12 µL with HyClone HyPure water H2O (Cytiva); negative controls included 1 nM pTXTL-P70a-T7rnap and myTXTL Sigma 70 Master Mix only. For each reaction, 10 µL was transferred to a PCR-clean polypropylene V bottom 96-well plate and covered with a silicone seal (Eppendorf, Mississauga, ON). Reactions were incubated at 29°C for 16 hours and final reactions stored at -20°C. To assess proteolytic activity, control (RNAP only) and PepO reactions were incorporated into fluorescent caseinase assays and collagenase assays. For caseinase assays, TXTL reactions were diluted 60-fold in TBS and 10 µL was added to each well. For type I and type IV collagenase assays, 10 µL of undiluted TXTL reactions were added to each well. Caseinase and type I collagenase assays were also conducted in the presence of 0.5 mM 1,10-phenanthroline. Mean fluorescence readings of the negative control (substrate with RNAP TXTL reaction) were subtracted from the experimental wells and relative fluorescence units (RFU) was plotted over time, with negative values adjusted to zero.

### Statistical Analyses

Statistics were performed in GraphPad Prism or Stata and graphs were prepared in GraphPad. GraphPad was used to assess data variance and normality (Shapiro-Wilk test) and statistical significance was evaluated as described in figure legends. All schematic illustrations were created with BioRender.com.

## Results

### Vaginal *Porphyromonas* species degrade type I collagen, type IV collagen and casein using secreted proteases

Given their phylogenetic relatedness to the periodontal pathogen, *P. gingivalis,* we sought to understand whether the two *Porphyromonas* species most frequently detected in the human vagina, *P. asaccharolytica* and *P. uenonis*, possess gingipain-like protease activities (14). Collagenase activity was evaluated using fluorescently quenched substrates (type I collagen or type IV collagen), where proteolytic digestion results in dequenching and measurable increases in fluorescence over time. Fluorometric collagenase assays confirmed that *P. asaccharolytica* and *P. uenonis* cell suspensions degrade type I collagen in a dose-dependent manner (Figure 1A). Next, collagenase activity was measured in cell-free supernatants from *P. asaccharolytica* and *P. uenonis.* These experiments validated that both organisms secrete proteases capable of degrading type I and type IV collagen (Figure 1B–C). Collagenase activity was further confirmed with gelatin zymography (Supplemental Figure 2), where *P. asaccharolytica* supernatants produced three distinct high molecular weight zones of clearing (∼85 kdA, 95 kDa, 120 kDa), while *P. uenonis* supernatants generated four separate zones of clearing in the gel: three high molecular weight (∼75 kDa, 90 kDa, 110 kDa) and one low molecular weight (30 kDa). Next, casein degradation was monitored to evaluate general proteolytic activity of *P. asaccharolytica* and *P. uenonis* secreted proteases. Due to its low-complexity tertiary structure, casein is regarded as a universal protease substrate that is highly susceptible to proteolytic degradation. Fluorometric assays revealed that supernatants from *P. asaccharolytica* and *P. uenonis* possess caseinase activity (Figure 2D). This was further confirmed using agar-based casein degradation assays, where *P. asaccharolytica, P. uenonis* and *P. gingivalis* cell suspensions all produced zones of clearing (Supplemental Figure 3). Although *P. asaccharolytica* and *P. uenonis* can degrade similar substrates to *P. gingivalis* (41), *P. gingivalis* showed substantially higher maximum fluorescence and area under the curve for collagen degradation from cell suspensions and secreted proteases (Supplemental Figure 4, Supplemental Table 1, Supplemental Table 2). The area under the curve for casein degradation by *P. gingivalis* was also much higher than for *P. asaccharolytica* or *P. uenonis,* while the maximum fluorescence and time to maximum fluorescence for casein degradation were comparable between all three *Porphyromonas* species (Supplemental Figure 4C, Supplemental Table 2). To understand if proteolytic activity might contribute to pathogenesis in the female genital tract, we evaluated whether a common commensal vaginal microbe is also capable of degrading collagen and casein. No collagenase or caseinase activity was detected from *Lactobacillus crispatus* cell suspensions in the fluorometric collagenase or caseinase assays (Supplemental Figure 5), suggesting that proteolytic activity could be a pathogenesis mechanism for opportunistic pathogens in the vaginal niche.

**Figure 1.**
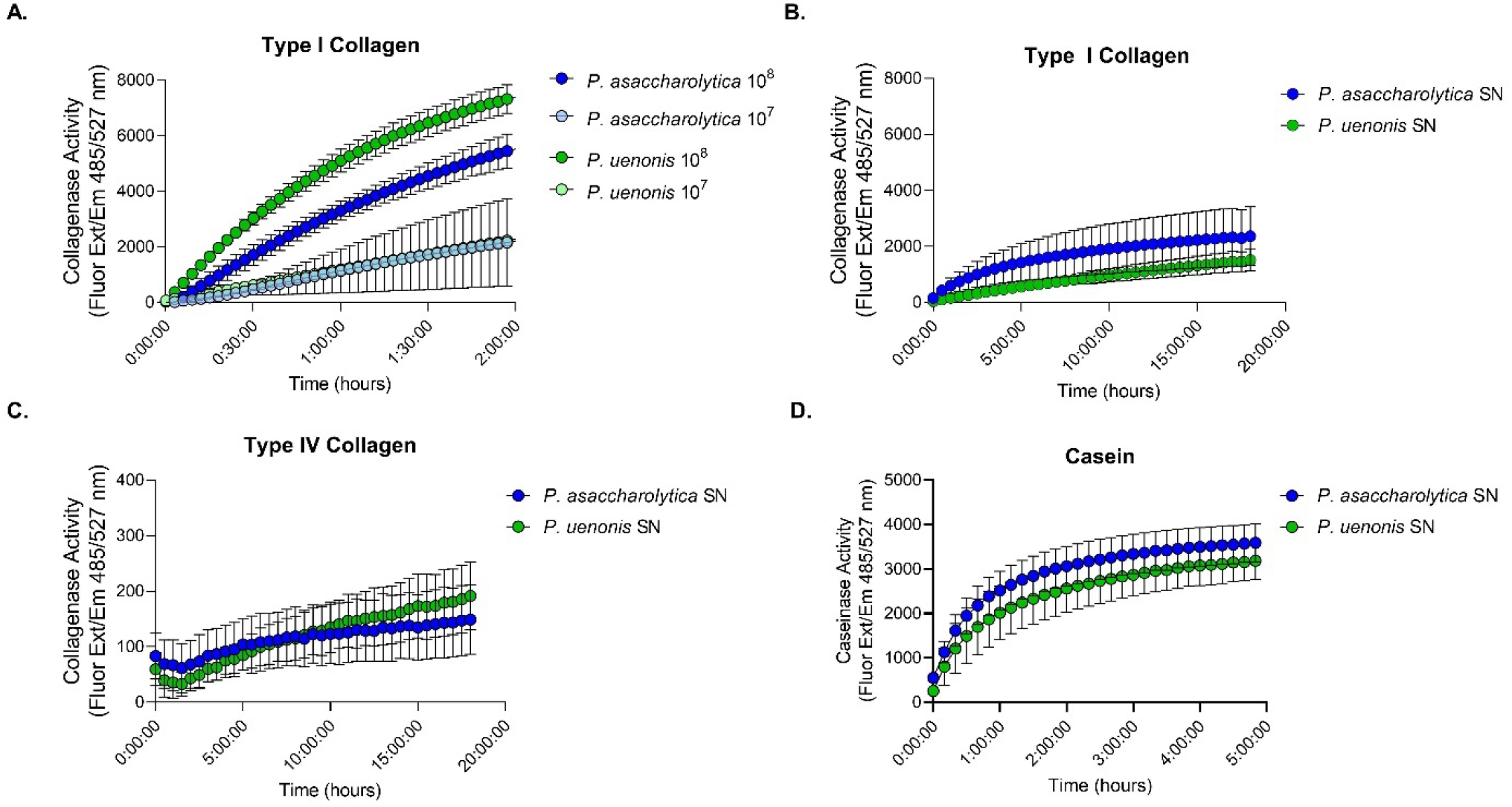
Proteolytic activity of vaginal *Porphyromonas* species. **(A)** Cell suspensions of *P. asaccharolytica* and *P. uenonis* at 10^7^ or 10^8^ cfu/reaction were incubated with fluorophore-conjugated type I collagen. Results are presented as mean ± standard error from three independent experiments performed in technical triplicate or quadruplicate. Collagen degradation was measured every three minutes by detecting the increase in fluorescence (Excitation 485 nm/Emission 527 nm) over a two- hour time course. **(B–C)** Cell-free supernatants of *P. asaccharolytica* and *P. uenonis* were incubated with fluorophore-conjugated **(B)** type I or **(C)** type IV collagen over an 18-hour time course; fluorescence was measured every 30 minutes. Results are presented as mean ± standard error from seven independent experiments (type I collagen) and four independent experiments (type IV collagen) performed in technical triplicate. **(D)** Cell-free supernatants of *P. asaccharolytica* and *P. uenonis* were incubated with fluorophore-conjugated casein over a five-hour time course; fluorescence was measured every ten minutes. Results are presented as mean ± standard error from five independent experiments performed in technical triplicate.

**Figure 2.**
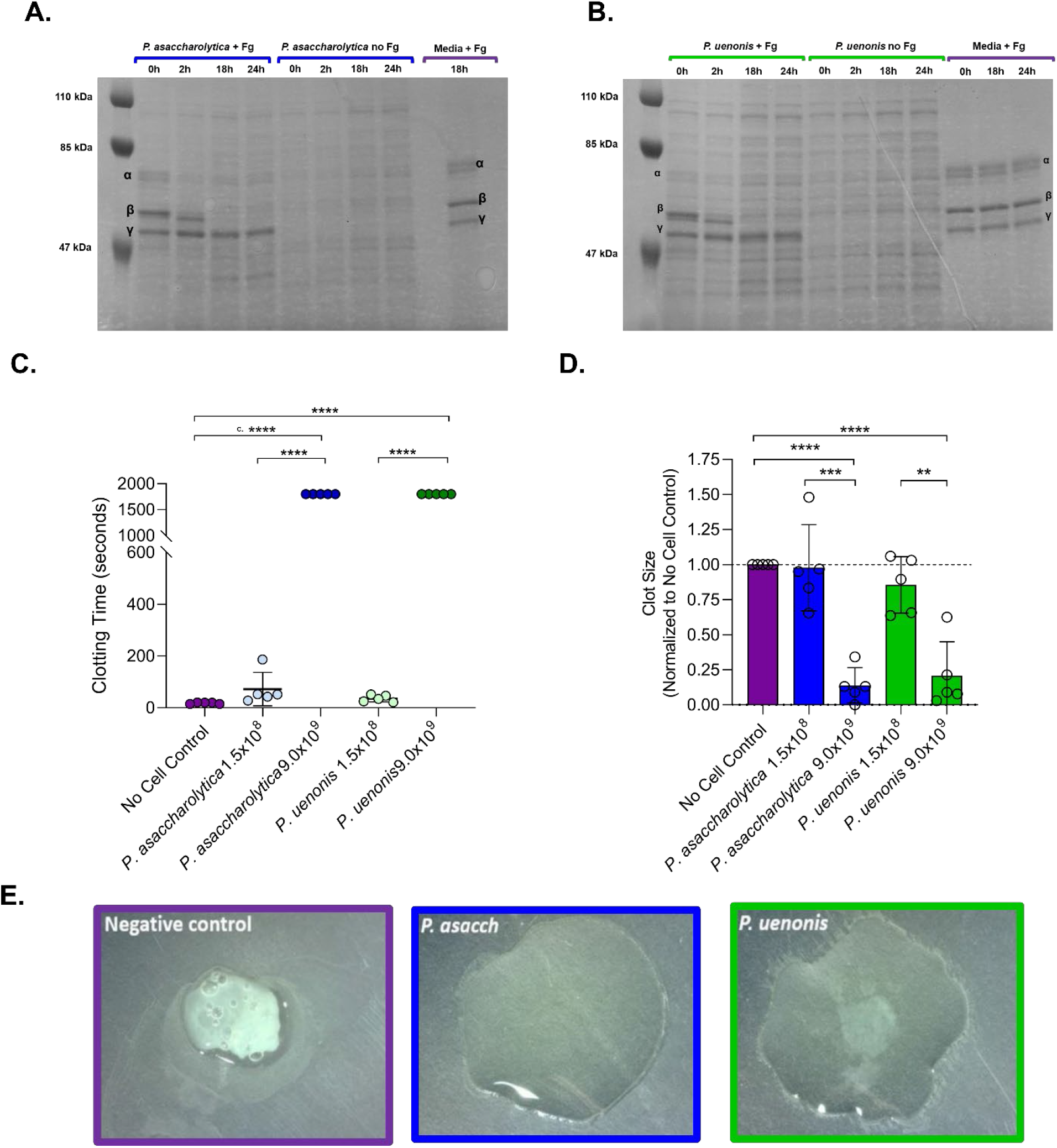
Vaginal *Porphyromonas* species degrade fibrinogen and impair clot formation. (A–B) SDS-PAGE of **(A)** *P. asaccharolytica* and **(B)** *P. uenonis* cell suspensions incubated with human fibrinogen (+Fg), saline (no Fg), or a cell-free media control (sBHI media +Fg). Samples collected over 24 hours were assessed for fibrinogen degradation, indicated by absence of bands corresponding to α, β, and γ fibrinogen chains in ‘*Porphyromonas* cells + Fg’ treatments compared to ‘no Fg’ or media controls. **(C)** Time from thrombin addition to fibrin clot formation. Citrated plasma was pre-incubated with cell suspensions of *P. asaccharolytica* or *P. uenonis* or no cell controls. Experiments were performed in technical duplicate and results are presented as mean ± standard error from five independent experiments. **(D)** Quantitative evaluation of clot size at the experimental endpoint (>30 minutes). Following clotting time assessment, samples were subjected to well area scans at absorbance 405 nm to indicate the average clot size. Experiments were performed in technical duplicate and control reactions without thrombin were used to blank the experimental reactions. Data were normalized to the average clot size of the no cell control within each experiment. Results are presented as mean ± standard error from five independent experiments. For clotting time, significance was assessed by one-way ANOVA with Holm-Sidak’s multiple comparisons test where **** p<0.0001. For endpoint clot size evaluation, significance was assessed by one-way ANOVA with Holm-Sidak’s multiple comparisons test where **** p<0.0001, *** p<0.0003, ** p=0.003 **(E)** Endpoint qualitative evaluation of fibrin clots (>30 minutes) after clotting time assay with cell suspensions of *P. asaccharolytica* (2.4×10^9^ cfu/reaction), *P. uenonis* (3.0×10^9^ cfu/reaction) or no cell control.

### *P. asaccharolytica* and *P. uenonis* inhibit fibrin clot formation through fibrinogen degradation

Since gingipains are known to degrade fibrinogen and exacerbate gum bleeding (41), we next investigated whether *P. asaccharolytica* and *P. uenonis* proteases can degrade fibrinogen and impair fibrin clot formation. To evaluate direct fibrinogen degradation, cell-free media controls and cell suspensions of *P. asaccharolytica* or *P. uenonis* were incubated in the presence and absence of human fibrinogen over a 24-hour time course. Samples removed at defined intervals were separated by SDS-PAGE and stained with Coomassie Brilliant Blue. In cell-free media controls, the fibrinogen α, β and γ chains remained intact throughout the experiment and, as expected, fibrinogen chains were absent from ‘*P. asaccharolytica* no Fg’ and ‘*P. uenonis* no Fg’ controls (Figure 2A–B). When *P. asaccharolytica* or *P. uenonis* were incubated with fibrinogen, complete degradation of the fibrinogen α and β chains was observed after 18 hours, while the γ chain remained intact (Figure 2A–B). To determine whether fibrinogen degradation translates to impaired fibrin clotting, thrombin-induced fibrin clot formation was measured after *Porphyromonas* cell suspensions were pre-incubated with citrated plasma. A significant delay in clot formation was observed with the highest dose of *P. asaccharolytica* or *P. uenonis* (9.0×10^9^ cfu/reaction) compared to the no cell control (Figure 2C, **** p<0.0001) or a lower dose of *P. asaccharolytica* or *P. uenonis* (Figure 2C; **** p<0.0001 vs. 1.5×10^8^). Turbidimetry at the experimental endpoint (30 minutes) allowed for quantitative evaluation of final clot size. In keeping with clotting times from Figure 2C, fibrin clot size was significantly reduced in samples exposed to *P. asaccharolytica* or *P. uenonis* at 9.0×10^9^ cfu/reaction compared with the no cell control (Figure 2D **** p<0.0001). For *P. asaccharolytica,* a significant reduction in clot size was observed in samples treated with 9.0×10^9^ cfu/reaction when compared to 1.5×10^8^ cfu/reaction (Figure 2D, *** p<0.0003). Similarily, clot sizes in samples treated with the highest dose of *P. uenonis* (9.0×10^9^ cfu/reaction) were significantly smaller than samples treated with a lower doses of *P. uenonis* (Figure 2D vs. 1.5×10^8^ cfu/reaction ** p=0.003). These findings were further confirmed by visual assessment of clot formation at assay endpoints (Figure 2E). Taken together, these results show that *P. asaccharolytica* and *P. uenonis* impair clot formation through proteolytic degradation of fibrinogen.

### Vaginal *Porphyromonas* species do not encode gingipain orthologs

Given the gingipain-like proteolytic activity observed in *P. asaccharolytica* and *P. uenonis* (Figures 1–2), we sought to confirm an earlier report that these organisms do not encode gingipain orthologs (78). Using BLASTP, gingipain protein sequences (RgpA, RgpB, Kgp) were queried against *P. asaccharolytica* and *P. uenonis* genomes (Supplemental Figure 6A), yielding zero hits in *P. asaccharolytica* and two hits in each *P. uenonis* 23387 genome (IMG Genome ID 2585427891 and 2528311143); of note, these *P. uenonis* genomes were not queried in the previous analysis (78). No BLASTP hits were identified in *P. uenonis* 60-3 as previously reported (78). One hit resulted from the Kgp BLAST search only (Supplemental Table 3; L215DRAFT_00230/JCM13868DRAFT_00677; 27% sequence identity) and another hit resulting from both the Kgp and RgpA BLAST queries (Supplemental Table 3; L215DRAFT_00128/JCM13868DRAFT_01423; 26–27% sequence identity). We further considered whether secreted proteases in *P. asaccharolytica* and *P. uenonis* share protein domains with gingipains. Evaluation of protein families (Pfams) in the gingipains revealed four conserved protein family (Pfam) domains, with RgpB containing one additional Pfam not found in RgpA or Kgp (Supplemental Table 4). All five Pfams were searched against all available genomes of *P. asaccharolytica* and *P. uenonis* to identify proteins containing gingipain Pfams (Supplemental Figure 6A). The peptidase C25 family (PF01364) search resulted in one hit in each of the genomes queried, while the cleaved adhesin domain (PF07675) returned hits in all *P. uenonis* strains, but none in *P. asaccharolytica* (Supplemental Table 4). To determine whether any hits from the BLAST search (Supplemental Table 3) and Pfam search (Supplemental Table 4) are gingipain orthologs, we performed a reciprocal BLASTP search against *P. gingivalis* (Supplemental Figure 6A). The C25 peptidase-containing proteins identified in the Pfam search against *P. asaccharolytica* (Poras_0230) and *P. uenonis* (L215DRAFT_00971) displayed the highest percent identity with the gingipains. When these sequences were queried against *P. gingivalis* using BLASTP, PGN_0022 emerged as the only significant hit with 36-37% identity and 97% query coverage (Supplemental Table 5). Since PGN_0022 has been characterized as PorU, the type 9 secretion sortase enzyme in *P. gingivalis* (79), the C25 peptidase-containing proteins identified in *P. asaccharolytica* and *P. uenonis* are likely to function as PorU orthologs rather than gingipains. Querying the cleaved adhesin domain-containing proteins from *P. uenonis* against *P. gingivalis* revealed two uncharacterized proteins: PGN_1611 and PGN_1733 as the top BLASTP hits (Supplemental Table 5). The Lys-gingipain (Kgp) was identified as fourth hit from both *P. uenonis* cleaved adhesin domain-containing protein BLASTP searches (Supplemental Table 5), but the percent identity and coverage are restricted to the shared pfam. Furthermore, results from multiple sequence alignments with the *P. gingivalis* gingipains showed ≤20% identity for all *P. asaccharolytica* and *P. uenonis* sequences. In summary, since all BLASTP alignments were less than 200 residues, full length alignments of query and subject sequences were ≤20%, reciprocal BLASTP searches did not return *P. gingivalis* gingipains as top hits, and our Pfam search did not identify *P. asaccharolytica* or *P. uenonis* proteins that appear to be gingipains, we concluded that these vaginal *Porphyromonas* species do not encode gingipain orthologs.

### Identification of collagenase candidates in *P. asaccharolytica* and *P. uenonis*

To identify other candidate proteins that may be responsible for the collagenolytic activity of vaginal *Porphyromonas* species, we first queried the MEROPs database (65) uncovering 59 and 63 known and predicted peptidases from *P. asaccharolytica* and *P. uenonis,* respectively. To identify putative collagenase enzymes from this list, we cross-referenced these peptidases with a list of protein annotation identifiers from known and predicted microbial collagenases in the BRENDA enzyme information database (66). This approach shortened the candidate peptidase list to 18 enzymes in *P. asaccharolytica* and 14 in *P. uenonis*. InterPro scans of each putative collagenase revealed proteins likely to be involved in cell wall synthesis or export machinery and narrowed the list to ten peptidases in *P. asaccharolytica* and nine peptidases in *P. uenonis* (Supplemental Figure 6B). Factoring in similarity to characterized collagenases, additional domains identified in InterPro and the presence of secretion signals, a final short list of the seven most promising candidate peptidases in each organism was generated (Table 1). Intriguingly, each organisms’ candidate collagenases could be organized into four groups: Ig-containing serine proteases (2x), C10 cysteine proteases (2x), M13 metalloproteases (1x) and U32 proteases (2x) (Table 1). Multiple sequence alignments of candidate collagenase pairs for a given type within each strain revealed low sequence identity (29%–45% (Table 1)). Conversely, BLAST searches between species allowed for identification of orthologous protein pairs in *P. asaccharolytica* and *P. uenonis*, with sequence identity ranging from 67% up to 100% (Table 1). Taken together, this suggests that within-genome pairs of candidate collagenases are likely to have resulted from a gene duplication event prior to speciation of *P. asaccharolytica* and *P. uenonis*.

**Table 1.**
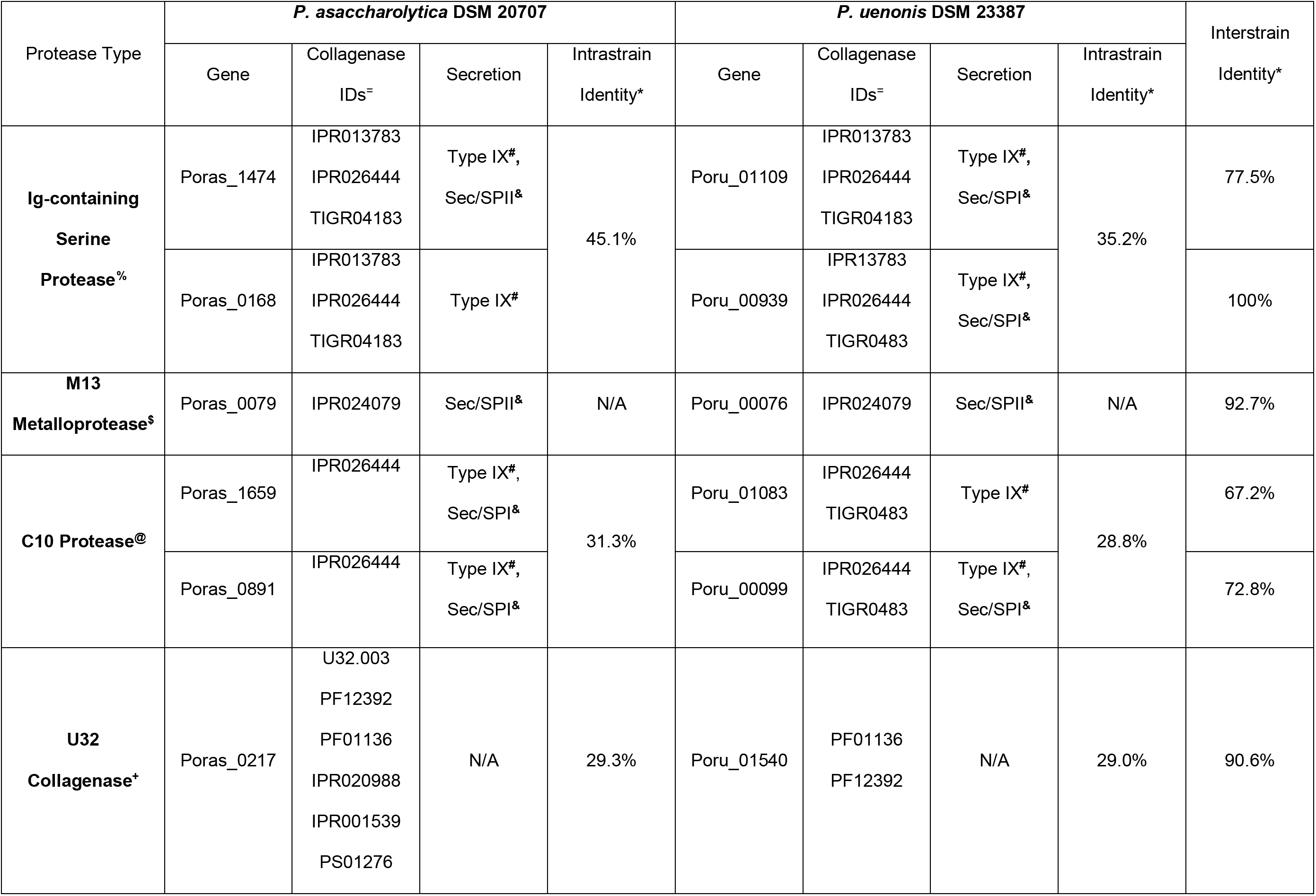
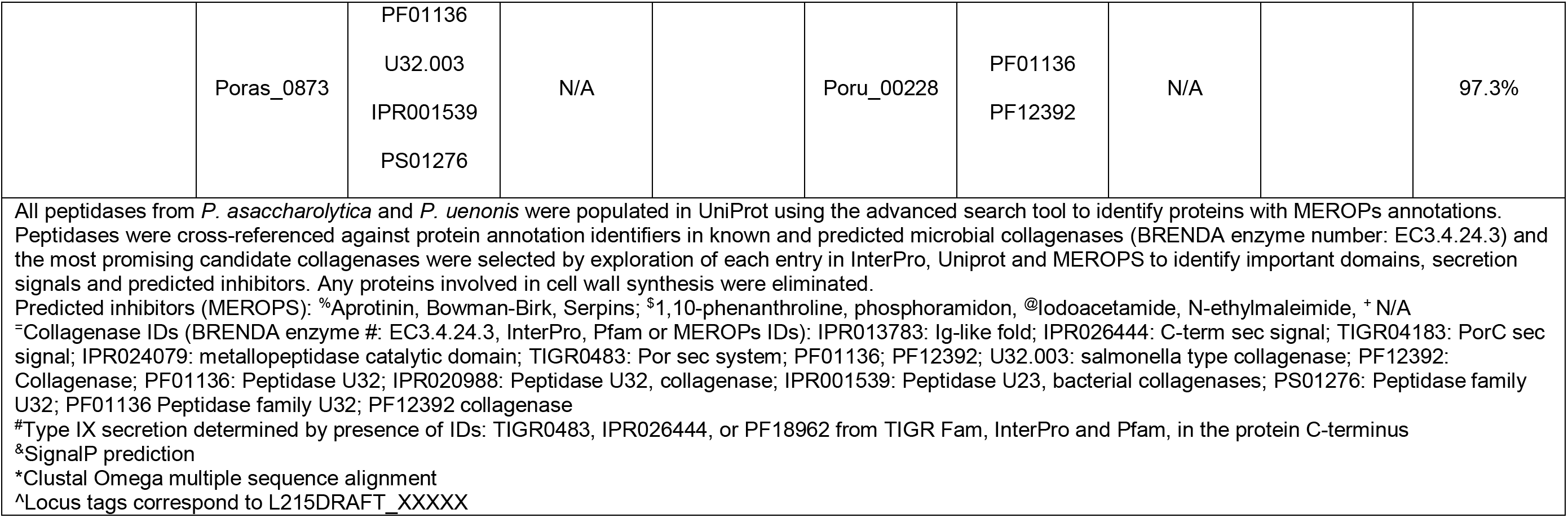
Candidate collagenases in P. asaccharolytica and P. uenonis.

Serine proteases within the short list all contained Ig-like folds (Table 1), which are also present in the binding domain of the well-characterized collagen degrading metalloproteases from *Clostridium histolyticum* (ColG, ColH) (80). The candidate cysteine proteases contained type 9 secretion system signal domains and a SpeB domain (Table 1), indicating sequence similarity with *Streptococcus pyogenes* streptopain, a cysteine protease capable of cleaving host components such as fibrinogen, immunoglobulins and complement proteins (81–83). The candidate metalloproteases from *P. asaccharolytica* (Poras_0079) and *P. uenonis* (Poru_00076) each contained a catalytic collagenase domain in addition to an M13 type metallopeptidase domain and a predicted N-terminal secretion signal (Table 1). Finally, two U32 collagenases were detected in each vaginal *Porphyromonas* species indicating an orthologous relationship with the *P. gingivalis* U32 collagenase PrtC (84, 85). Of note, none of these putative U32 collagenases were found to possess secretion signals indicative of localization to the extracellular space (Table 1).

### Vaginal *Porphyromonas* species encode metalloproteases targeting collagens, casein and fibrinogen

To narrow down candidate enzymes responsible for collagenolytic and fibrinogenolytic activities, inhibitors of the predicted serine, cysteine and metalloproteases (Table 1) were incorporated into functional assays. Cell-free supernatants from *P. asaccharolytica* and *P. uenonis* were incubated with type I collagen in the presence of three doses of 1,10-phenanthroline, iodoacetamide or aprotinin to inhibit metallo-, cysteine and serine proteases, respectively (Figure 3A**–**D, Supplemental Figure 6A**–**F). Treatment of *P. asaccharolytica* supernatants with 1,10-phenanthroline resulted in decreased collagenase activity, with a statistically significant reduction in max enzyme activity observed with the highest dose of inhibitor (Figure 3A**–**B; * 0.2 mM vs. 0.02 mM p=0.004; ** 0.2 mM vs. 0.002 mM p<0.0001, Supplemental Figure 6A). Iodoacetamide treatment revealed a trend toward a dose-dependent decrease in collagenase activity in individual experiments (Supplemental Figure 7), but this trend was not observed when multiple experiments were combined (Figure 3A**–**B, Supplemental Figure 6). However, when *P. asaccharolytica* supernatants were treated with a combination of 1,10- phenanthroline and iodoacetamide, there was a further reduction in collagenase activity compared to the 1,10-phenanthroline only treatment (Figure 3A**–**B, 1,10-Ph 0.2 mM vs. 1,10-Ph + Iodo * p=0.0378, Supplemental Figure 6). Although aprotinin treatment revealed a trend towards decreased activity in the time-course, there was no statistically significant decrease in max collagenase activity (Figure 3A**–**B, Supplemental Figure 6). For *P. uenonis,* 1,10-phenanthroline also inhibited collagenase activity (Figure 3C**–**D, Supplemental Figure 6), with a significant reduction in max collagenase activity (Figure 3D; 0.2 mM vs. 0.02 mM p=0.002; 0.2 mM vs. 0.002 mM p=0.0043). However, treatment with aprotinin or iodoacetamide did not reduce *P. uenonis* collagenase activity (Figure 3C**–**D, Supplemental Figure 6) and the combination treatment of 1,10-phenanthroline and iodoacetamide did not provide an additional reduction in collagenase activity or max fluorescence (Figure 3C**–**D) as observed in *P. asaccharolytica* (Figure 3A-B). Taken together, these results demonstrate that both *P. asaccharolytica* and *P. uenonis* possess secreted metalloproteases that coordinate type I collagen degradation, while *P. asaccharolytica* also appears to secrete a cysteine protease capable of degrading type I collagen.

**Figure 3.**
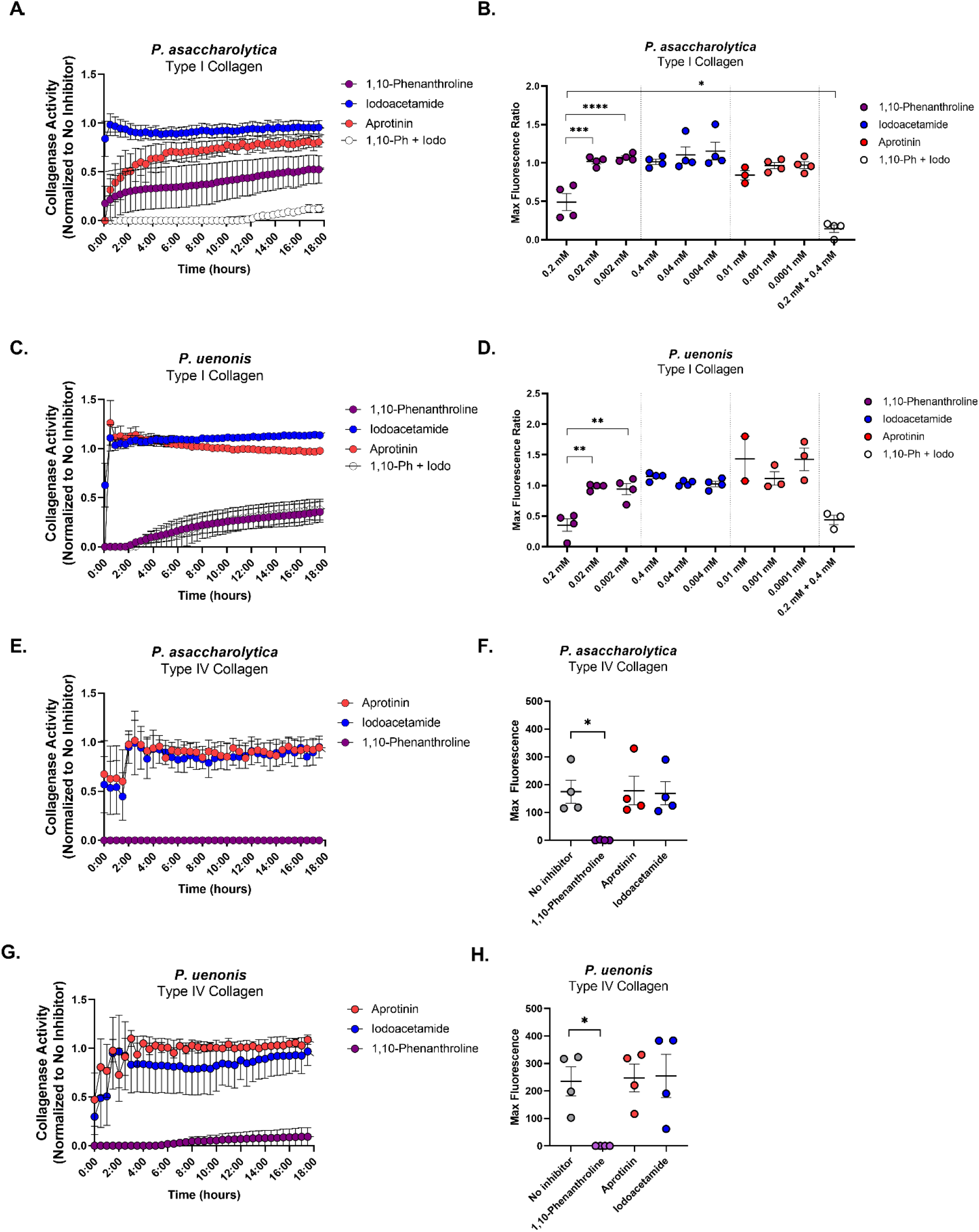
Vaginal *Porphyromonas* protease types displaying type I and type IV collagenase activity. Cell-free supernatants of **(A-B, E-F)** *P. asaccharolytica* and **(C-D, G-H)** *P. uenonis* were incubated with fluorophore-conjugated **(A-D)** type I collagen or **(E-H)** type IV collagen in the presence of the metalloprotease inhibitor 1,10-phenanthroline, the cysteine protease inhibitor, iodoacetamide, or the serine protease inhibitor, aprotinin. **(A,C)** Type I collagen degradation was measured every 30 minutes by detecting fluorescence (Excitation 485 nm/Emission 527 nm) over an 18-hour time course in the presence of 0.2 mM 1,10-phenanthroline, 0.4 mM iodoacetamide, 0.01 mM aprotinin or 0.2 mM 1,10-phenanthroline + 0.4 mM iodoacetamide. Results are presented as a ratio normalized to the no inhibitor control (set to 1.0). **(B)** Maximum fluorescence of *P. asaccharolytica* collagenase activity, expressed as a ratio normalized to the no inhibitor control, in the presence of three doses of the inhibitors. Results are presented as mean ± standard error from four independent experiments or three independent experiments (*P. asaccharolytica* + aprotinin 0.01 mM) performed in technical triplicate. Statistical significance was assessed with a one-way ANOVA and Tukey’s post-hoc comparison where 1,10-phenanthroline 0.2 mM vs. 0.02 mM **** p<0.0001, 1,10-phenanthroline 0.2 mM vs. 0.002 mM *** p=0.0004 and 1,10-phenanthroline 0.2 mM vs. 0.2 mM 1,10-phenanthroline + 0.4 mM iodoacetamide * p=0.0387. **(D)** Maximum fluorescence ratios of *P. uenonis* collagenase activity, expressed as a ratio normalized to the no inhibitor control, in the presence of three doses of the inhibitors. Results are presented as mean ± standard error from four independent experiments or two independent experiments (*P. uenonis* + aprotinin 0.01 mM) performed in technical triplicate. Statistical significance was assessed with a one-way ANOVA and Tukey’s post-hoc comparison, where 1,10-phenanthroline 0.2 mM vs. 0.02 mM ** p=0.002, 1,10-phenanthroline 0.2 mM vs. 0.002 mM ** p=0.0043. **(E-H)** Type IV collagen degradation was measured every 30 minutes by detecting fluorescence (Excitation 485 nm/Emission 527 nm) over an 18-hour time course in the presence of 0.2 mM 1,10-phenanthroline, 0.4 mM iodoacetamide, 0.01 mM aprotinin or 0.2 mM 1,10-phenanthroline + 0.4 mM iodoacetamide. Results are presented as a ratio normalized to the no inhibitor control and presented as mean ± standard error from three independent experiments performed in technical triplicate. **(F)** Maximum fluorescence of *P. asaccharolytica* supernatant collagenase activity in the presence of inhibitors. Results are presented as mean ± standard error from four independent experiments performed in technical triplicate. Statistical significance was assessed with a one-way ANOVA and Tukey’s post-hoc comparison where 1,10-phenanthroline 0.2 mM vs. no inhibitor * p=0.0352. **(H)** Maximum fluorescence of *P. uenonis* supernatant collagenase activity in the presence of inhibitors. Results are presented as mean ± standard error from four independent experiments performed in technical triplicate. Statistical significance was assessed with a one-way ANOVA and Tukey’s post-hoc comparison where 1,10- phenanthroline 0.2 mM vs. no inhibitor * p=0.0398.

Next, we evaluated whether the same protease classes degrade type IV collagen and casein. For both *Porphyromonas* species, the metalloprotease inhibitor 1,10-phenanthroline completely abrogated type IV collagenase activity from supernatants, while serine and cysteine protease inhibitors did not significantly reduce activity (Figure 3E–H). *P. asaccharolytica* caseinase activity was similar to its type I collagenase activity (Figure 4A–B) as proteolytic activity was reduced by treatment with 1,10- phenanthroline, and treatment with both 1,10-phenanthroline and iodoacetamide produced an additional downward shift in enzyme activity that resulted in a significant reduction in max fluorescence when compared to the no inhibitor control (Figure 4A, C; p=0.0413). In keeping with the type I collagen and type IV collagen degradation results, *P. uenonis* caseinase activity was significantly decreased by treatment with the metalloprotease inhibitor 1,10-phenanthroline (Figure 4B,D no inhibitor vs. 1,10- phenanthroline p=0.018), while aprotinin and iodoacetamide treatment did not inhibit activity (Figure 4B,D). Further to this, the 1,10-phenanthroline/iodoacetamide combination did not offer any additional reduction in protease activity (Figure 4B,D).

**Figure 4.**
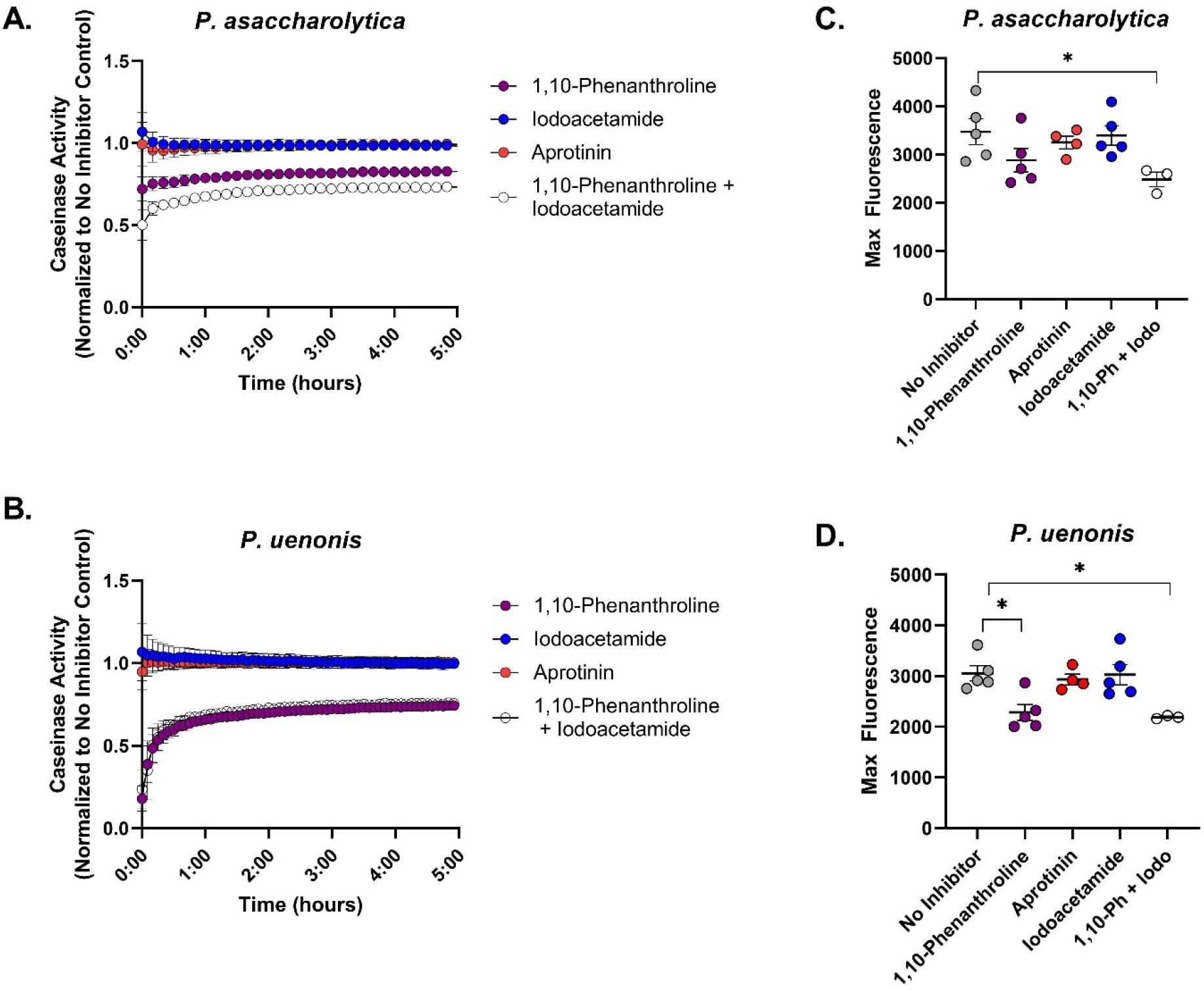
*P. asaccharolytica* and *P. uenonis* secreted metalloproteases have broad substrate specificity. (A-B) Cell-free supernatants of **(A)** *P. asaccharolytica* and **(B)** *P. uenonis* were incubated with fluorophore-conjugated casein in the presence of the metalloprotease inhibitor 1,10-phenanthroline (0.2 mM), the cysteine protease inhibitor iodoacetamide (0.4 mM), or the serine protease inhibitor aprotinin (0.01 mM). Casein degradation was measured every 10 minutes by detecting the increase in fluorescence (Excitation 485 nm/Emission 527 nm) over a 5-hour time course. Results are presented as a ratio normalized to the no inhibitor control and presented as mean ± standard error from five independent experiments performed in technical triplicate. **(C)** Maximum fluorescence of *P. asaccharolytica* supernatant caseinase activity in the presence of inhibitors. Results are presented as mean ± standard error from five independent experiments performed in technical triplicate. Statistical significance was assessed with a one-way ANOVA and Tukey’s post-hoc comparison, where inhibitor combination vs. no inhibitor * p=0.0413. **(D)** Maximum fluorescence of *P. uenonis* supernatant caseinase activity in the presence of inhibitors. Results are presented as mean ± standard error from five independent experiments performed in technical triplicate. Statistical significance was assessed with a one-way ANOVA and Tukey’s post-hoc comparison, where 1,10-phenanthroline 0.2 mM vs. no inhibitor p=0.0145, inhibitor combination vs. no inhibitor p=0.0180.

Inhibitors were also incorporated into fibrinogen degradation assay to determine whether fibrinogen is proteolyzed by the same enzyme classes as those observed in our collagen and casein experiments. When *P. asaccharolytica* supernatants were incubated with fibrinogen, complete degradation of the fibrinogen α and β chains was observed after 48 hours (Figure 5A, no inhibitor), while the γ chain remained intact, producing the same degradation profile observed in experiments with cell suspensions (Figure 2A). Delayed fibrinogen degradation by *P. asaccharolytica* was observed in the presence of 1,10-phenanthroline, but not with aprotinin or iodoacetamide. This suggests that secreted metalloproteases from *P. asaccharolytica* contribute to fibrinogen degradation (Figure 5A). Fibrinogen degradation patterns from *P. uenonis* supernatants differed from the results obtained with cell suspensions. While the fibrinogen α chain was degraded after two hours, the β and γ chains remained intact for the 48-hour time-course (Figure 5B). This finding suggests that *P. uenonis* may degrade fibrinogen using both secreted and cell surface-associated proteases. Interestingly, the secreted *P. uenonis* protease that contributes to degradation of the fibrinogen α chain is not impacted by the inhibitors included in this study (Figure 5B), implying that the secreted fibrinogenolytic enzyme from *P. uenonis* is distinct from the secreted collagen and casein degrading enzymes.

**Figure 5.**
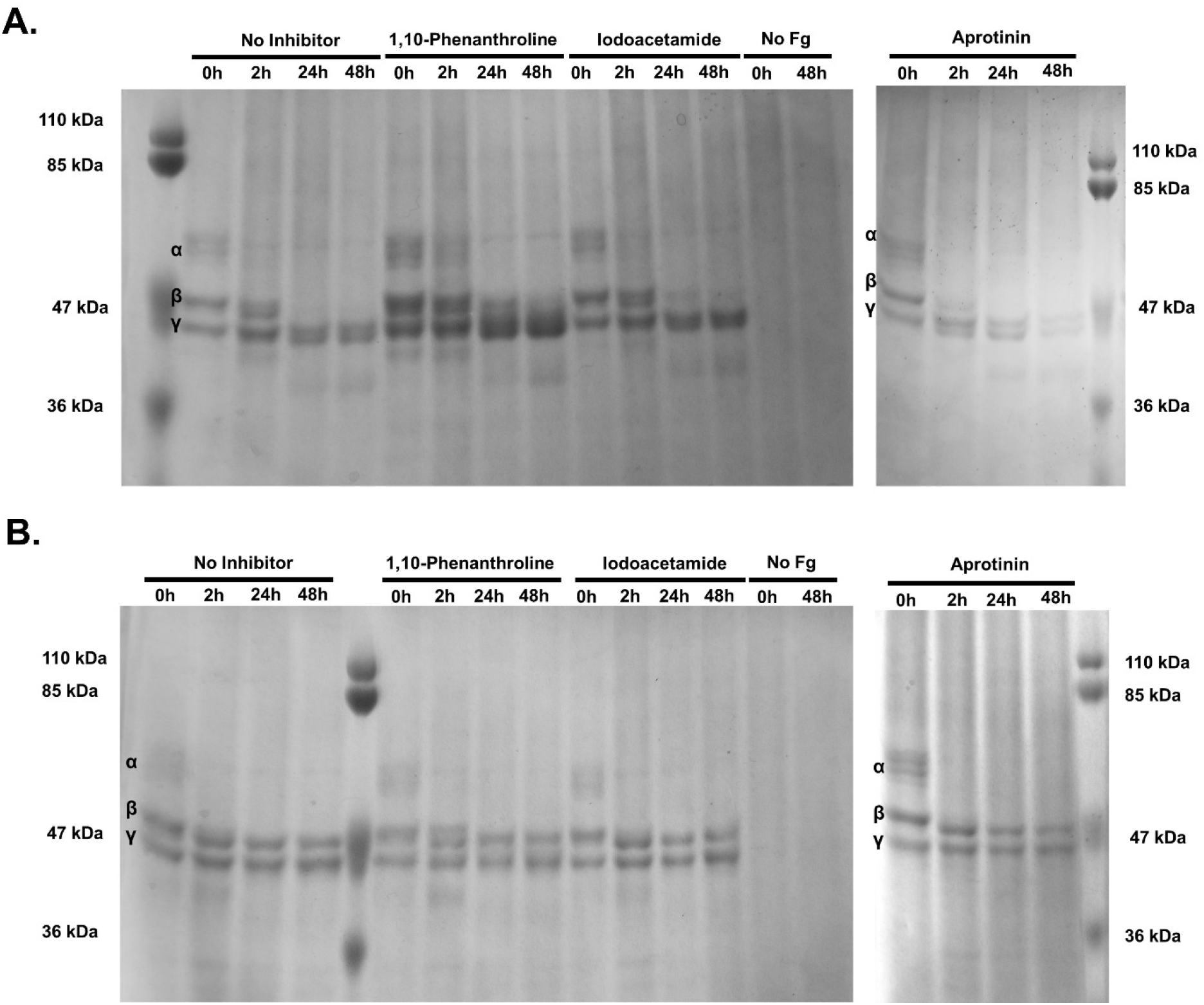
*P. asaccharolytica* metalloproteases degrade fibrinogen. **(A)** SDS-PAGE of *P. asaccharolytica* supernatants incubated with human fibrinogen in the presence of 1,10-phenanthroline (0.5 mM), iodoacetamide (1mM) or aprotinin (0.01 mM) compared to no inhibitor and no fibrinogen controls. Samples collected over 48 hours were assessed for fibrinogen degradation, indicated by absence of bands corresponding to α, β, and γ fibrinogen chains as denoted within gel images. **(B)** SDS-PAGE of *P. uenonis* supernatants incubated with human fibrinogen in the presence of 1,10- phenanthroline (0.5 mM), iodoacetamide (1mM) or aprotinin (0.01 mM) compared to no inhibitor and no fibrinogen controls. Samples collected over 48 hours were assessed for fibrinogen degradation, indicated by absence of bands corresponding to α, β, and γ fibrinogen chains.

Finally, we sought to further characterize the *Porphyromonas* M13 metalloproteases identified in our bioinformatics inquiries (Table 1; Figure 6A). Exploration of other *Porphyromonas* species detected in the urogenital tract and commonly isolated from other human body sites as well, revealed that the M13 metalloproteases are ubiquitous (Figure 6B). Notably, the M13 metalloprotease in *P. gingivalis* has been previously characterized as PepO, a secreted endopeptidase involved in host attachment/invasion and proteolytic activation of endothelin, a potent peptide that induces vasoconstriction (62, 63, 86). To determine whether the vaginal *Porphyromonas* M13 metalloproteases can coordinate collagenase and caseinase activity, Poras_0079 (*pepO)*, was cloned and expressed in myTXTL *in vitro* transcription/translation system. Expression of PepO was confirmed via SDS-PAGE with the appearance of a 76 kDa protein in PepO reactions, that was absent in the control RNA polymerase only reactions (Figure 7A). Fluorescent protease assays revealed that PepO is capable of degrading casein and type I collagen, but not type IV collagen (Figure 7B–D). Further, the metalloprotease inhibitor, 1,10-phenanthroline, fully abrogated type I collagenase and caseinase activity of PepO (Figure 7B–C).

**Figure 6.**
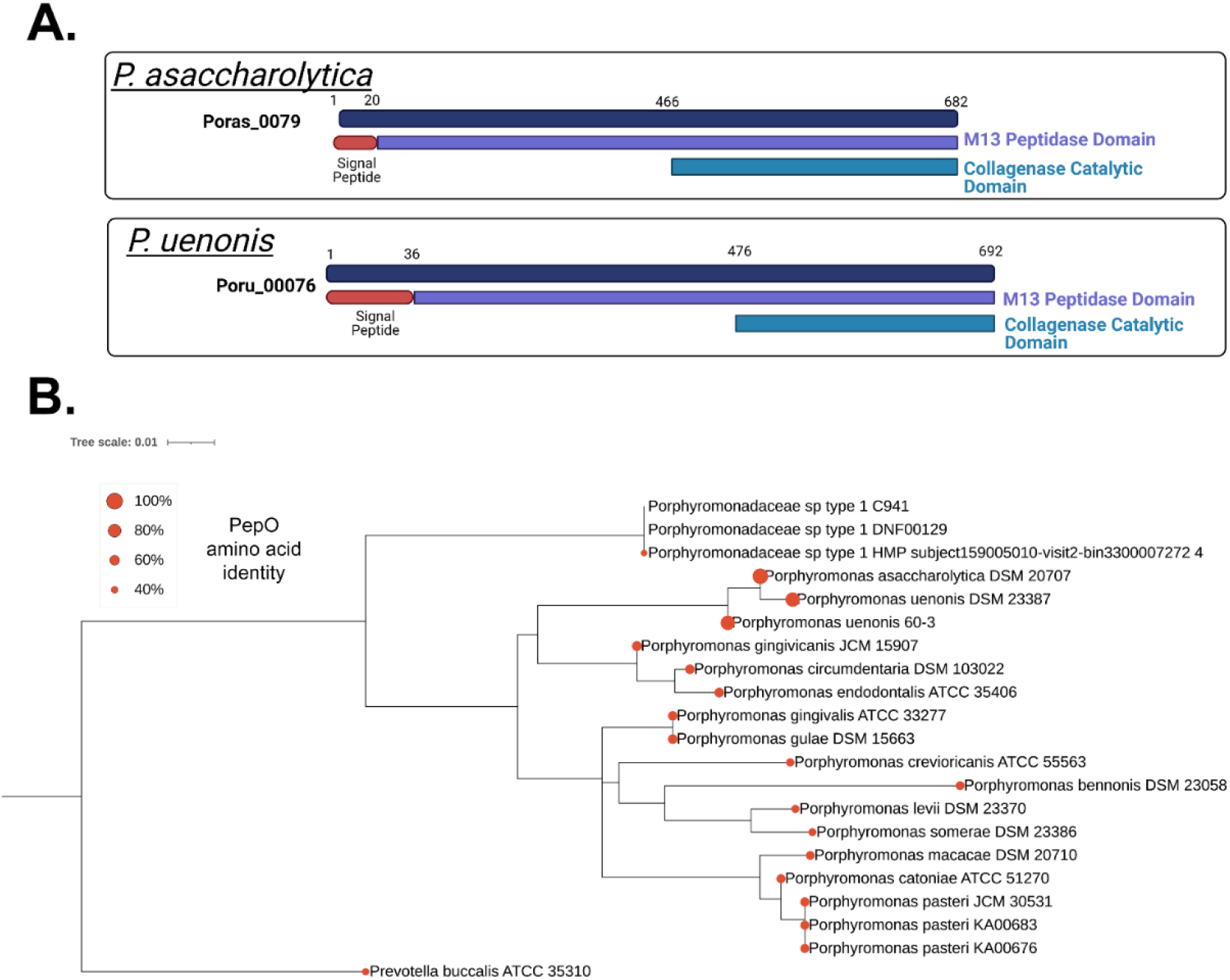
Secreted PepO metallopeptidases identified in *P. asaccharolytica* and *P. uenonis*. **(A)** Domain structure of candidate host-degrading PepO metalloproteases from *P. asaccharolytica* and *P. uenonis*. **(B)** 16S rRNA gene phylogeny of *Porphyromonas* species identified in the urogenital tract. The Maximum Likelihood tree was created using RAxML and rooted to *Prevotella buccalis* (same order, different family). Red circles on each leaf indicate the percent amino acid identity of that species’ PepO ortholog compared to *P. asaccharolytica* PepO (Poras_0079 = 100%). The taxon previously identified as uncultivated *Porphyromonas* species type 1 was found to encompass two cultured, but unsequenced isolates (DNF00129 and C941, >99% 16S rRNA gene identity over 1120 nt). We also identified a metagenome assembled genome (MAG) in IMG/MER that encoded a 958 bp 16S rRNA gene fragment >99.5% identical to those from the cultured isolates DNF00129 and C941. Since this MAG represents the only genome sequence available for this species, we used it in our PepO queries, identifying an orthologue 42% identical to *P. asaccharolytica* PepO. Since this taxon’s PepO ortholog was more distantly related to *P. asaccharolytica* PepO than the PepO ortholog identified in *Prevotella buccalis* (47% identical), it may represent a novel genus or family; we therefore chose to label this taxon *Porphyromonadaceae* sp. type 1. Our phylogenetic analysis, along with inquiries through the Genome Taxonomy Database, furthermore indicated that vaginal isolates KA00683 and KA00676 should be designated as belonging to the species *P. pasteri*, with each containing a PepO ortholog 58– 59% identical to that from *P. asaccharolytica*.

**Figure 7.**
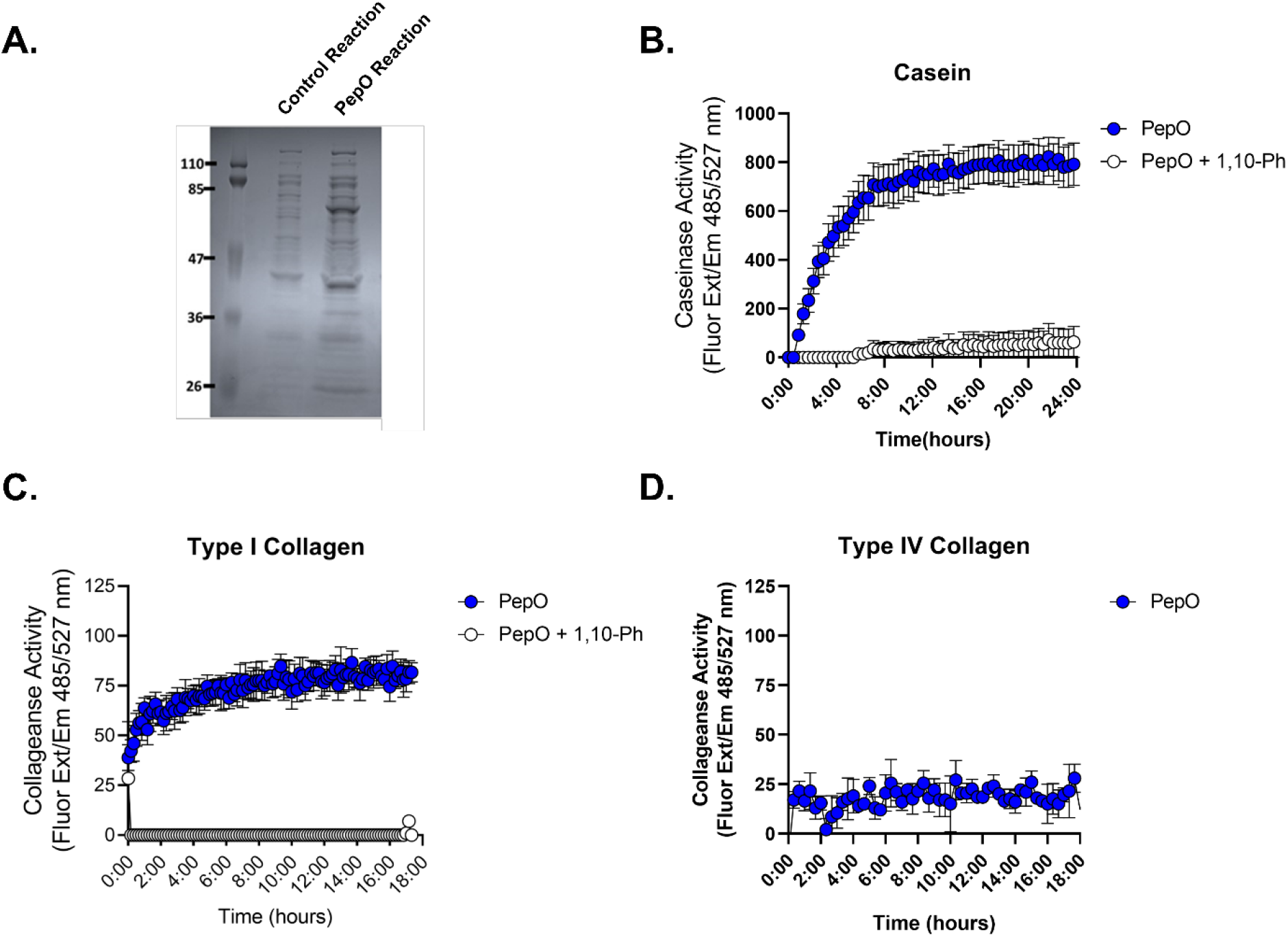
The *P. asaccharolytica* metalloprotease PepO degrades casein and type I collagen **(A)** SDS- PAGE of control *in vitro* transcription/translation (TXTL) reaction (RNA polymerase only) and PepO TXTL reaction. **(B-D)** PepO TXTL reactions were incubated with **(B)** FITC-casein or **(C)** fluorophore- conjugated type I collagen in the presence of 0.5 mM 1,10-phenantrholine. Casein degradation was measured every 30 minutes by detecting the increase in fluorescence (Excitation 485 nm/Emission 527 nm) over a 24-hour time course and results are presented as mean ± standard error from three independent experiments. Collagen degradation was measured every 10 minutes by detecting the increase in fluorescence (Excitation 485 nm/Emission 527 nm) over an 18-hour time course. Results are presented as mean ± standard error from five independent experiments. **(D)** PepO TXTL reactions were incubated with type IV collagen over an 18-hour time course and results are presented as mean ± standard deviation from one independent experiment.

## Discussion

It is well established that mucinase activity is elevated during BV (87–90), and these activities have been attributed to *Gardnerella* (89, 91) and *Prevotella* species (89, 92, 93). In support of this, sialidase activity has been utilized as a diagnostic marker for BV (92, 94–97). Although most studies have focused on degradation of mucin glycans, proteolytic activity in vaginal fluid has also been linked with BV status (56–58) and described among predominant BV-associated bacteria (59, 60). Isolates of *Prevotella bivia* from women with preterm premature rupture of membranes (PPROM) were shown to secrete proteases that degrade elastin, collagen, casein and gelatin (59). In another study screening bacterial strains from women with PPROM, preterm labour or puerperal infection, protease activity was confirmed in phylogenetically diverse Gram-negative and Gram-positive organisms. This included the BV-associated bacteria *Gardnerella vaginalis*, *Prevotella bivia* (formerly *Bacteroides bivius)* and *P. asaccharolytica* (formerly *Bacteroides asaccharolyticus*) (60). The authors found that *P. bivia* and *P. asaccharolytica* exhibited collagen and casein degradation, while *G. vaginalis* exclusively degraded casein. Our findings confirm the casein/collagen degradation capacity of *P. asaccharolytica* and extend these functional activities to *P. uenonis.* Furthermore, we demonstrate that both vaginal *Porphyromonas* species also degrade type IV collagen, commonly found in reproductive tissues (98, 99), and provide the first description of the diverse protease types capable of coordinating these activities. To understand whether proteolytic activity is also observed among commensal bacteria that inhabit the vaginal niche, we investigated proteolytic activity of *Lactobacillus crispatus*, demonstrating that *L. crispatus* is not capable of degrading type I collagen or casein. The absence of detectable proteolytic activity from a commensal *Lactobacillus* strain and the growing evidence for secreted proteolytic activity from BV-associated bacteria (59, 60), including vaginal *Porphyromonas* species, suggests that degradation of host proteins could be an important virulence trait of opportunistic pathogens in the female genital tract.

Our results demonstrate that vaginal *Porphyromonas* species are capable of directly degrading fibrinogen and impairing fibrin clot formation. Fibrinogen is detected in vaginal lavage fluid (100, 101) and is targeted by other reproductive pathogens (102, 103). Although the implications of altered fibrinogen levels in the female reproductive tract are not clear, impaired clotting functions could have severe consequences during labour and postpartum. Fibrinogen is known to increase substantially during pregnancy (104, 105), and decreased plasma fibrinogen levels have been associated with increased severity of postpartum haemorrhage (106). Further to this, genetic fibrinogen abnormalities are significantly associated with miscarriage, placental abruption and postpartum hemorrhage (107–110).

Within the female genital tract, collagens are found within the vagina, cervix, uterus and pelvic floor and their composition and content is significantly altered throughout pregnancy and during labour (111). Vaginal and cervical tissue is primarily composed of fibrillar type I and III collagens, with type I collagen playing a critical role in tissue integrity (112–114). Type IV collagen is typically found within basement membranes and is enriched within the placenta (98). Both type I and type IV collagens are found within chorioamniotic membranes at the maternal-fetal interface (99). During pregnancy, cervical collagens (type I, III) undergo a shift toward increased solubility and decreased abundance, contributing to cervical softening (115–117), while increased collagenase activity is observed during cervical ripening to prepare for dilation and parturition (118, 119). Importantly, cervical remodelling during term and preterm labour occurs via the same mechanisms with host matrix metalloproteinases (MMPs) coordinating cervical collagen degradation (120). Premature preterm rupture of the membrane (PPROM) has been associated with infection (121, 122), increased host MMP collagenase activity (123, 124) and decreased collagen content (125). Furthermore, microbial collagenases can reduce the tensile strength of chorioamniotic membranes *ex vivo* (126), and collagenase activity has been detected in clinical isolates from PPROM patients (59, 60). Taken together, these findings imply that host and microbial modulation of collagen within the cervix and chorioamniotic membranes could play critical roles in preterm labour and PPROM. In the present study, our findings indicate that *P. asaccharolytica* and *P. uenonis* secreted proteases are capable of degrading both type I and type IV collagens, uncovering a possible mechanism for how these microbes contribute to the initiation of preterm labour.

*P. asaccharolytica* and *P. uenonis* are phylogenetically related to *P. gingivalis,* an opportunistic pathogen that drives periodontal disease (30), disseminates through the bloodstream (35) and contributes to adverse pregnancy outcomes (34, 127). Many of these outcomes are driven by the *P. gingivalis* gingipains, cysteine proteases that degrade host extracellular matrix components and immune factors (41). Our investigations demonstrate that *P. asaccharolytica* and *P. uenonis* possess gingipain-like activities, including degradation of type I and type IV collagen, casein and fibrinogen. However, key differences in total enzyme activity were observed. *P. gingivalis* supernatants possessed higher total collagenase and caseinase enzyme activity than the vaginal *Porphyromonas* species based on the maximum RFU, time to max RFU and area under the curve values. Previous comparative genomic analyses of *Porphyromonas* species included two *P. asaccharolytica* genomes (DSM 20707, PR426713P-I) and one *P. uenonis* genome (60–3), and used a reciprocal BLAST (BLASTall) to confirm that *P. asaccharolytica* and *P. uenonis* do not encode gingipain orthologs (78). Expanding on these findings, our study used both BLAST and domain (Pfam) queries, and included all available *P. asaccharolytica* and *P. uenonis* genomes including two additional *P. uenonis* genomes (DSM 23387; IMG Genome IDs 2528311143 and 2585427891) that were not included in the previous study. In agreement with the previous study (78), our findings did not return any gingipain ortholog hits from reciprocal BLASTP or Pfam searches. This absence of gingipain orthologs is in keeping with the observed differences in functional proteolytic activity. Taken together, these results prompted an investigation of novel candidate proteases.

Our bioinformatics inquiries identified five candidate secreted collagenases in each vaginal *Porphyromonas* species. By incorporating protease inhibitors into our functional assays, we determined that *Porphyromonas* secreted metalloproteases degrade collagens (type I, IV), casein and fibrinogen. For *P. uenonis,* we observed consistent results with metalloprotease inhibitor 1,10-phenanthroline exclusively inhibiting degradation with casein and collagens (type I,IV), while no inhibitors blocked fibrinogen degradation. Importantly, no additional reduction in activity was observed when the cysteine protease inhibitor iodoacetamide was included with 1,10-phenanthroline in type I collagenase or caseinase assays, suggesting that secreted metalloproteases are solely responsible for degradation of these substrates by *P. uenonis*. With *P. asaccharolytica*, on the other hand, we observed that iodoacetamide caused a dose-dependent reduction in *P. asaccharolytica* type I collagenase activity. Furthermore, when 1,10-phenanthroline and iodoacetamide were combined, there was an additional decrease in *P. asaccharolytica* type I collagenase and caseinase activity relative to treatment with 1,10- phenanthroline alone. These results suggest that although *P. asaccharolytica* and *P. uenonis* possess the same candidate collagenases, only *P. asaccharolytica* appears to secrete both metallo and cysteine proteases in the experimental conditions used in our study. This is further supported by collagen zymogram results, where banding patterns revealed different sizes of collagenases from *P. asaccharolytica* and *P. uenonis.* Future investigations will need to address whether pH or redox state may affect the activity of the cysteine proteases. Additionally, due to the general proteolytic activity of the metallo and cysteine proteases, it is plausible that secreted proteases may degrade other proteins in the supernatants, including other proteases.

Our bioinformatics approach highlighted the U32 collagenases as likely candidates to confer proteolytic activity, however, we were unable to find any sequence-based evidence for secretion to the extracellular space or localization to the cell surface in *P. asaccharolytica* or *P. uenonis*. As such, these enzymes are unlikely to be found in culture supernatants. The collagenase activity of the *P. gingivalis* U32 collagenase, PrtC, has been confirmed (84, 85, 128), but these studies used cell lysates (85), recombinant PrtC (84) or heterologous expression systems (128). To our knowledge, no studies have determined the subcellular localization of PrtC, and sequence analysis does not reveal any indication of secretion signals. In fact, additional studies have confirmed that the majority of collagenase activity from *P. gingivalis* is attributed to the gingipains (129, 130). On the other hand, characterized U32 collagenases from other bacteria are known to be secreted (131, 132). The active site residues, protease type and inhibitors of U32 collagenase remain unknown, but the lack of evidence for secretion in *Porphyromonas* species makes these U32 collagenases improbable candidates for the proteolytic activity observed in this study.

Since 1,10-phenanthroline inhibited the secreted proteolytic activity of *P. asaccharolytica* and *P. uenonis*, the M13 metalloproteases were expected to confer the observed protease activity. These predicted collagenases from *P. asaccharolytica* and *P. uenonis* each contain an N-terminal signal sequence, M13 peptidase domain and a catalytic collagenase domain. The *P. asaccharolytica* M13 metalloprotease was cloned and expressed in an *in vitro* transcription/translation system and confirmed to contribute type I collagenase and caseinase activity. Phylogenetic analysis of the *P. asaccharolytica* and *P. uenonis* M13 metalloproteases revealed an orthologous relationship with the *P. gingivalis* endopeptidase, PepO (PgPepO), and confirmed the presence of PepO orthologs in other vaginal and oral *Porphyromonas* species. Previous work characterizing *P. gingivalis* PepO revealed sequence conservation with the human endothelin converting enzyme 1 (ECE-1), which proteolytically processes inactive endothelin (big endothelin) into active endothelin. Activated endothelin peptides can induce vasoconstriction and cellular proliferation, alter vascular permeability and activate inflammatory cells (133, 134). PgPepO was confirmed to possess ECE-1 like activity, converting all three subtypes of big endothelin to active endothelin (62). Numerous studies have also demonstrated that PgPepO plays a role in cellular invasion and intracellular survival of *P. gingivalis* (63, 86). However additional substrates for this endopeptidase and the functional consequences of bacterial endothelin activation have yet to be explored.

PepO orthologs have also been characterized in select *Lactobacillus* species: *Lactobacillus lactis* and *Lactobacillus rhamnosus* PepO can proteolyze casein, but these enzymes are either confirmed or predicted to localize in the cytoplasm (135, 136). PepO has also been explored in *Streptococcus* species, including *Streptococcus pneumoniae* and *Streptococcus pyogenes* (Group A Streptococci; GAS). In *S. pneumoniae,* PepO is detected on the bacterial cell surface and in culture supernatants. *S. pneumoniae* PepO binds to host cells and fibronectin, facilitating bacterial adhesion and invasion (137). Streptococcal PepO also interacts with the host immune system, but there is conflicting evidence on whether this contributes to immune evasion or activation. While some studies demonstrate that *S. pneumoniae* PepO binds plasminogen and complement proteins (C1q), contributing to escape from fibrin clots and complement attack (137, 138), others demonstrate that PepO enhances macrophage autophagy and bactericidal activity via TL2/4 activation (139, 140). Surprisingly, none of these studies have shown any proteolytic activity of *S. pneumoniae* PepO. In GAS, PepO has been detected both in the cytoplasm and as a secreted protein (141, 142). Similar to *S. pneumoniae*, GAS PepO contributes to complement evasion via C1q binding (142). GAS PepO participates in regulating quorum sensing via direct degradation of peptide pheromones secreted by GAS (141). Taken together, this body of work highlights the broad substrate specificity and diverse functionality of bacterial PepO. To our knowledge, our work characterizing *P. asaccharolytica* PepO is the first to demonstrate degradation of host extracellular matrix (type I collagen), adding another substrate to the proteolytic repertoire of PepO. The findings that PepO can degrade regulatory proteins secreted by host cells (*P. gingivalis* PepO: endothelin) and bacterial cells (GAS PepO: quorum sensing peptides) suggests that PepO enzymes could play a role in the dysregulation of proteolytic cascades. Further investigation is needed to better understand the substrates targeted by PepO proteins in complex mucosal body sites such as the female reproductive tract to reveal how this enzyme contributes to the pathogenesis of phylogenetically diverse bacteria.

## Acknowledgements

The authors would like to thank Dr. Antoine Dufour and Dr. Morley Hollenberg for technical assistance and advice on this work.

