## Supplemental Data for "Protease activities of vaginal *Porphyromonas* species disrupt coagulation and extracellular matrix in the cervicovaginal niche"

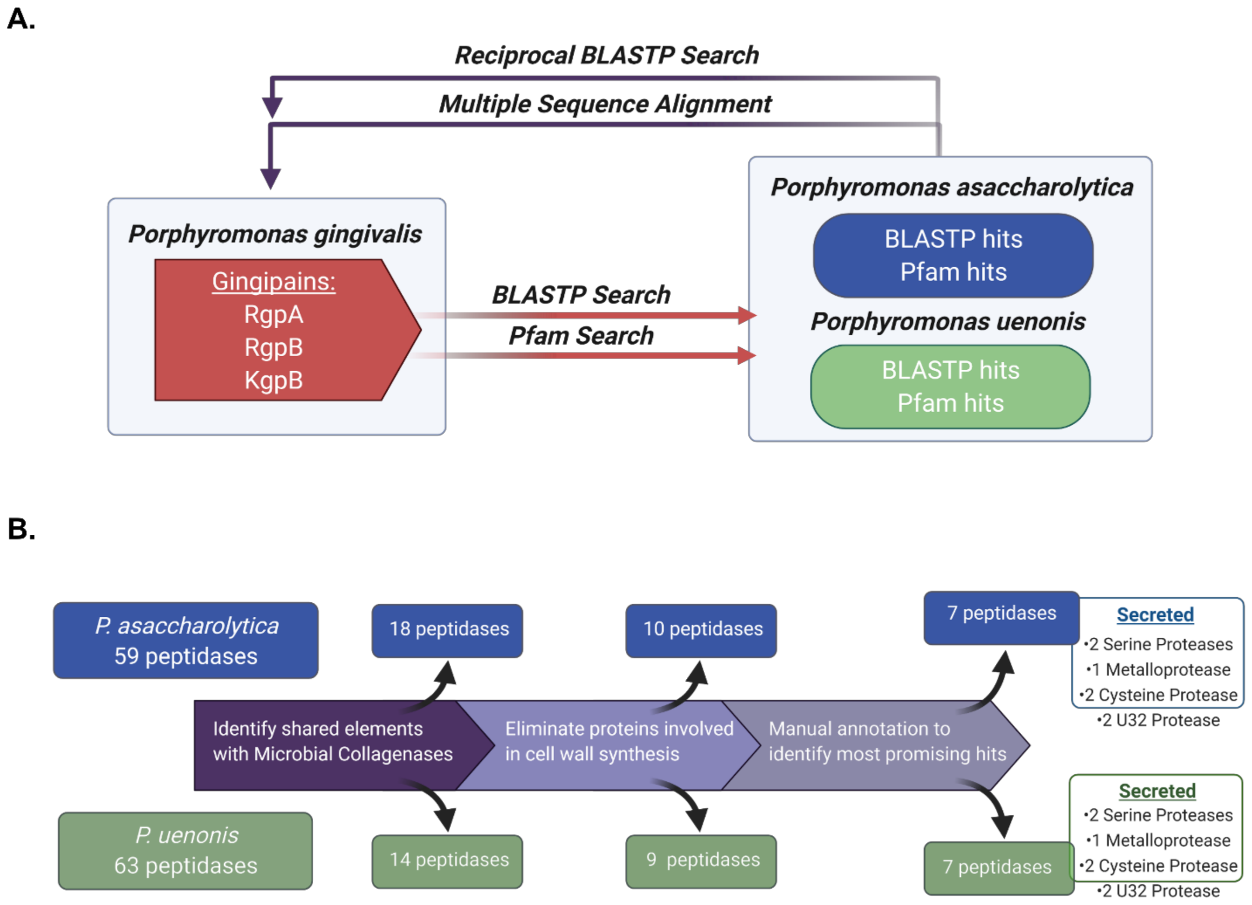


**Supplemental Figure 1. Schematic of bioinformatics approaches. (A)** Gingipain amino acid sequences for RgpA, RgpB and KgpB were used in BLASTP and Pfam searches against *P. asaccharolytica* and *P. uenonis* to search for gingipain orthologs. Resulting hits were subject to a reciprocal BLASTP search against *P. gingivalis* and a multiple sequence alignment with each gingipain. **(B)** Identification of candidate collagenases in *P. asaccharolytica* and *P. uenonis.* The MEROPS database was searched for all peptidases in *P. asaccharolytica and P. uenonis. Porphyromonas* peptidases were cross-referenced against protein annotation identifiers found in known and predicted microbial collagenases (BRENDA enzyme number: EC3.4.24.3). Candidate collagenases were explored in InterPro and UniProt to eliminate proteins involved in cell wall synthesis and identify the most promising hits. Images were prepared using BioRender.com**.**

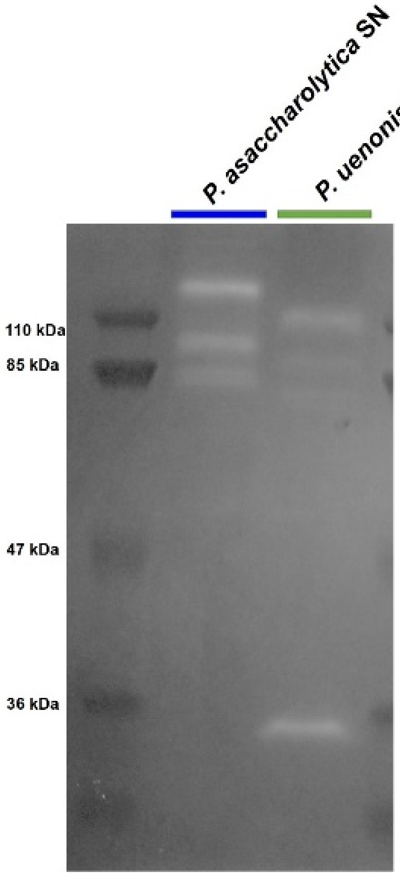


**Supplemental Figure 2. Gelatin zymography of *P. asaccharolytica* and *P. uenonis* cell-free supernantants.** Non-denaturing SDS-PAGE gelatin (type I collagen) zymogram loaded with 40 µg per well of *P. asaccharolytica* or *P. uenonis* supernatant. Zones of clearing observed in Coomassie-stained gels after gel renaturing and enzyme activation. Representative zymogram from three independent experiments.

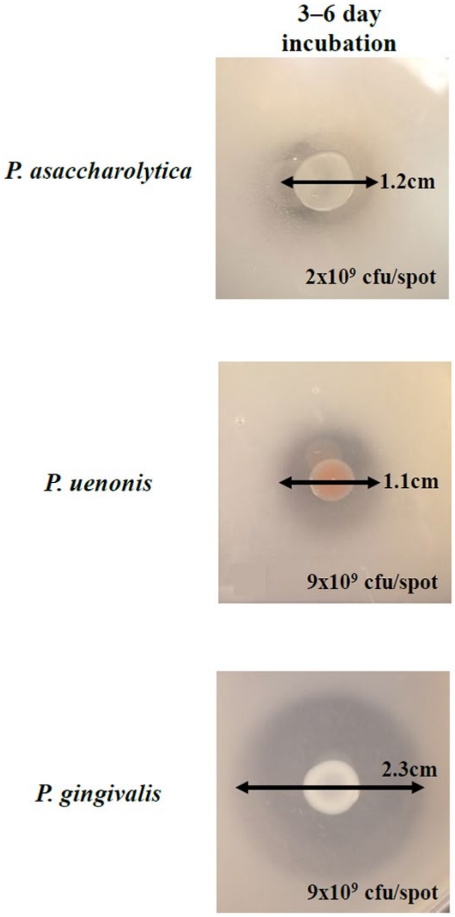


**Supplemental Figure 3. Casein degradation ability of *P. asaccharolytica, P. uenonis* and *P. gingivalis.*** Bacterial cell suspensions were spotted onto casein agar plates. Zones of clearing indicating casein degradation were measured after incubation for 3 or 6 days at designated doses of bacteria.

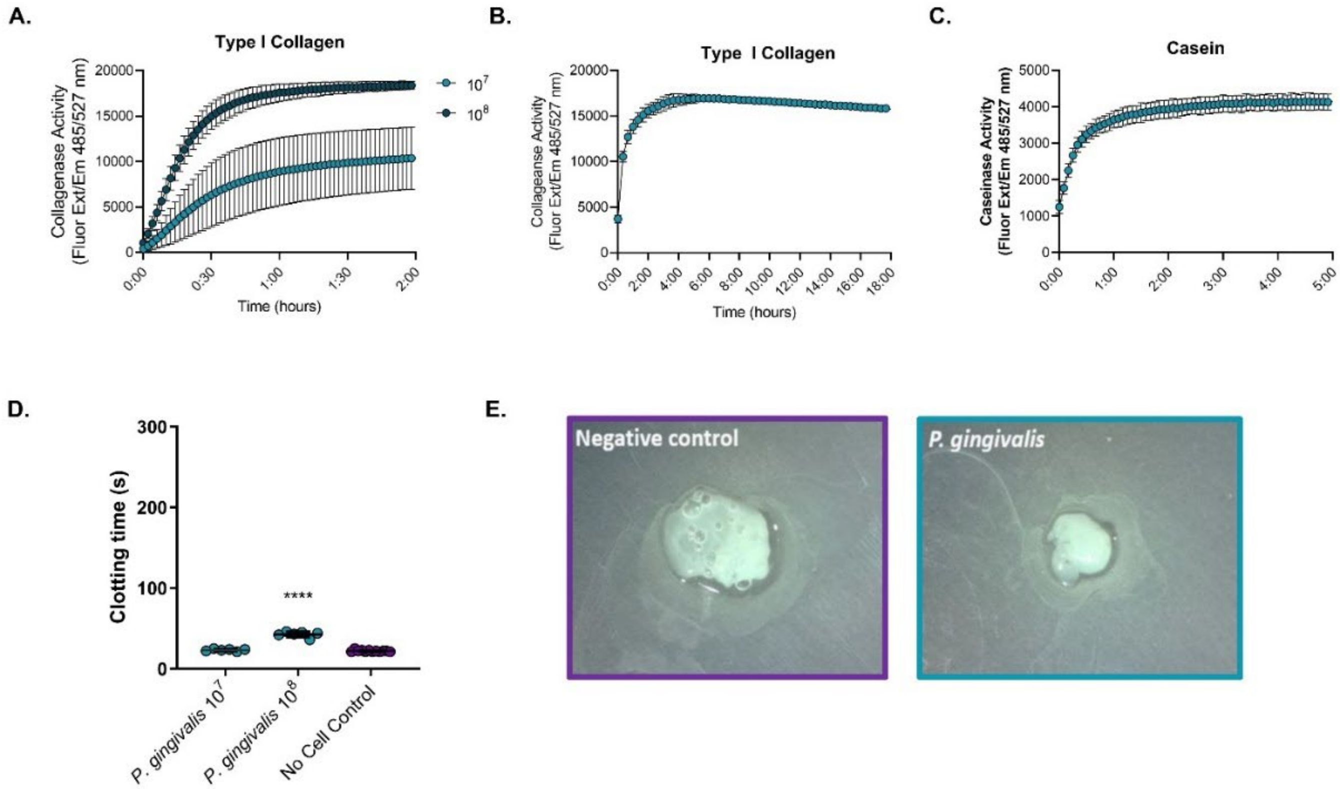


**Supplemental Figure 4. Protease activity of *P. gingivalis.*** **(A)** Cell suspensions of *P. gingivalis* at $10^7$ or $10^8$ cfu/reaction were incubated with fluorophore-conjugated type I collagen. Results are presented as mean ± standard error from two independent experiments performed in technical triplicate or quadruplicate. Collagen degradation was measured every three minutes by detecting the increase in fluorescence (Excitation 485 nm/Emission 527 nm) over a two-hour time course. **(B)** Cell-free supernatants of *P. gingivalis* were incubated with fluorophore-conjugated type I collagen over an 18-hour time course; fluorescence was measured every 30 minutes. Results are presented as mean ± standard error from two independent experiments performed in technical triplicate. **(C)** Cell-free supernatants of *P. gingivalis* were incubated with fluorophore-conjugated casein over a five-hour time course; fluorescence was measured every ten minutes. Results are presented as mean ± standard error from two independent experiments performed in technical triplicate. **(D)** Time to fibrin clot formation after thrombin addition following pre-incubation of citrated plasma with no cell controls or cell suspensions of *P. gingivalis* at $10^7$ or $10^8$ cfu/reaction. Experiments were performed in technical duplicate and results are presented as mean ± SEM from two independent experiments. **(E)** Endpoint qualitative evaluation of fibrin clots (>30 minutes) after clotting time assay from no cell control or *P. gingivalis* cell suspension ($10^8$ cfu/mL).

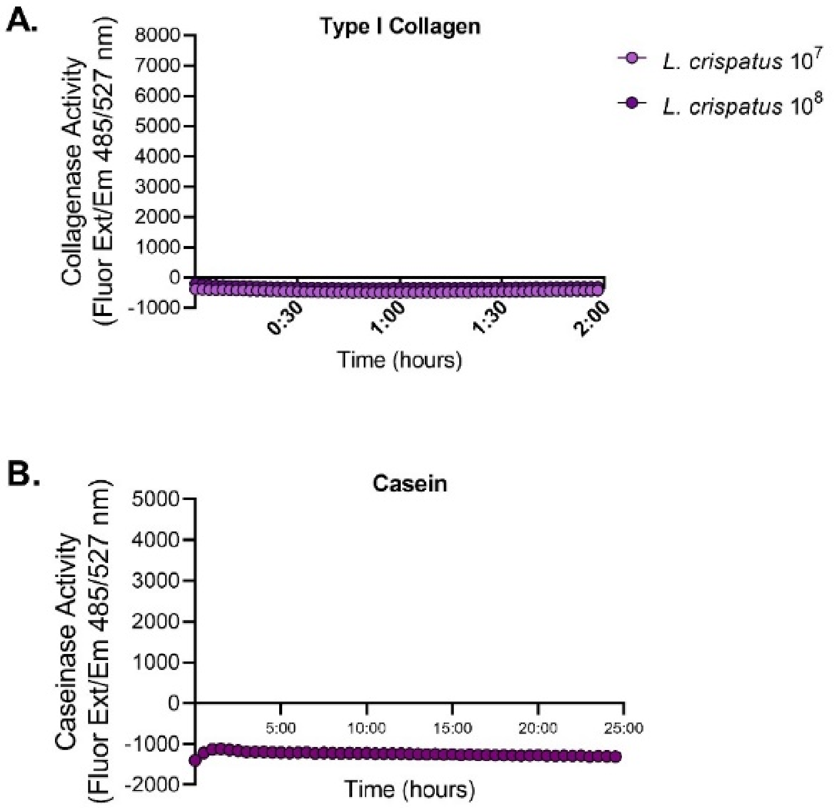


**Supplemental Figure 5. *Lactobacillus crispatus* does not degrade type I collagen or casein. (A)** Cell suspensions of *L. crispatus* at $10^7$ or $10^8$ cfu/reaction were incubated with fluorophore-conjugated type I collagen. Collagen degradation was measured every 3 minutes by detecting the increase in fluorescence (Excitation 485 nm/Emission 527 nm) over a two-hour time course. Results are presented as mean ± standard deviation from one independent experiment performed in technical quadruplicate. **(B)** Cell suspensions of *L. crispatus* at $3x10^6$ cfu/reaction were incubated with fluorophore-conjugated casein. Casein degradation was measured every 30 minutes by detecting the increase in fluorescence (Excitation 485 nm/Emission 527 nm) over a 24-hour time course. Results are presented as mean ± standard deviation from one independent experiment performed in technical triplicate.

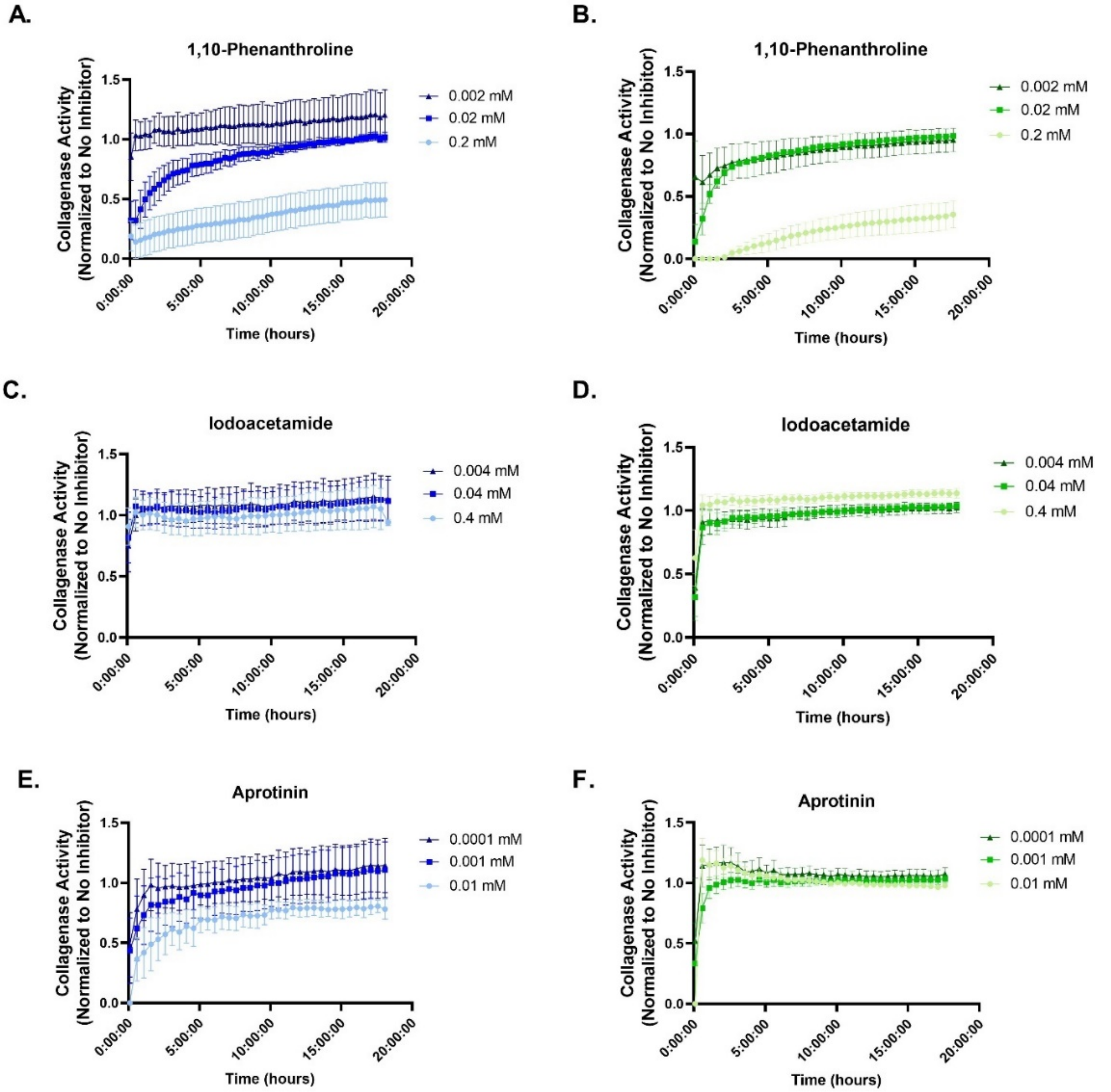


**Supplemental Figure 6. Dose response inhibition of *P. asaccharolytica* and *P. uenonis* type I collagenase activity with protease inhibitors.** Cell-free supernatants of **(A,C,E)** *P. asaccharolytica* (blue) or **(B,D,F)** *P. uenonis* (green) were incubated with fluorophore-conjugated type I collagen in the presence of three different doses of **(A-B)** the metalloprotease inhibitor 1,10-phenanthroline, **(C-D)** the cysteine protease inhibitor iodoacetamide, or **(E-F)** the serine protease inhibitor aprotinin. Collagen degradation was measured every 30 minutes by detecting the increase in fluorescence (Excitation 485 nm/Emission 527 nm) over an 18-hour time course. Results are presented as a ratio normalized to the no inhibitor control and presented as mean ± standard error from four independent experiments or two independent experiments (aprotinin 0.01 mM dose) performed in technical triplicate.

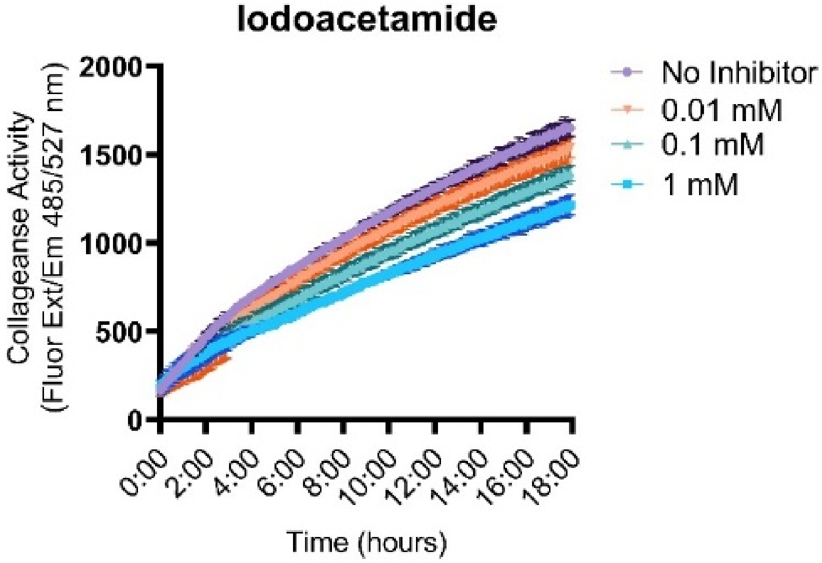


**Supplemental Figure 7. Dose-dependent inhibition of *P. asaccharolytica* type I collagenase activity with the cysteine protease inhibitor iodoacetamide.** *P. asaccharolytica* cell-free supernatant was incubated with fluorophore-conjugated type I collagen in the no inhibitor control or in presence of three different doses of the cysteine protease inhibitor iodoacetamide. Collagen degradation was measured every five minutes by detecting the increase in fluorescence (Excitation 485 nm/Emission 527 nm) over an 18-hour time course. Results are presented as mean ± standard deviation from one independent experiment.

**Supplemental Table 1. Type I collagenae activity of *Porphyromonas* species and *Lactobacillus crispatus* cell suspensions.**

| | **$10^7$ cfu/reaction** | | | | **$10^8$ cfu/reaction** | | | |
|---|---|---|---|---|---|---|---|---|
| | Max RFU | Time to Max RFU | Slope | AUC | Max RFU | Time to Max RFU | Slope | AUC |
| *P. asaccharolytica* | 2204.7 ± 1593.5 | 120 ± 0 | 1181 ± 119.1 | 2204 ± 207.7 | 5526 ± 628.2 | 120 ± 0 | 2857 ± 55.1 | 6170 ± 91.1 |
| *P. uenonis* | 2281 ± 168.7 | 120 ± 0 | 1111 ± 10.2 | 2349 ± 17.7 | 7389 ± 515.8 | 120 ± 0 | 3556 ± 62.7 | 9260 ± 86.74 |
| *P. gingivalis* | 10418 | 120 ± 0 | 4388 ± 439 | 15454 ± 746.3 | 18427 | 120 ± 0 | 6030 ± 316 | 30981± 184.3 |
| *L. crispatus* | -392 | 0 | -12 | 0 | -201 | 0 | -24 | 0 |

RFU = relative fluorescence units

AUC = area under the curve

**Supplemental Table 2. Secreted collagenase and caseinase activity of *Porphyromonas* species.**

| | Type I Collagen | | | | Casein | | | |
|---|---|---|---|---|---|---|---|---|
| | Max RFU | Time to Max RFU | Slope | AUC | Max RFU | Time to Max RFU | Slope | AUC |
| *P. asaccharolytica* | 2396.8 ± 1542.4 | 1091 ± 1.7 | 99.8 ± 9.5 | 30991 ± 1742 | 3442 + 262.3 | 291 + 1.9 | 424 ± 24.8 | 13717 ± 270.7 |
| *P. uenonis* | 1059.5 ± 392.0 | 1093 ± 1.0 | 79.3 ± 2.8 | 15700 ± 552.4 | 3010.5 + 204.5 | 291.3 + 2.4 | 422.2 ± 21.9 | 11532 ± 209.3 |
| *P. gingivalis* | 17024 ± 373 | 335 ± 62.6 | 89 ± 11 | 292886 ± 413.9 | 4153 ± 236.6 | 265 ± 17.3 | 283 ± 22 | 78127 ± 202.8 |

RFU = relative fluorescence units

AUC = area under the curve

**Supplemental Table 3. Gingipain BLASTP hits in *P. uenonis* DSM 23387.**

| Gingipain Query | BLASTP Hits[#] | E-value | % Sequence Identity | Blast Hit Coverage (residues) | Subject Length |
|---|---|---|---|---|---|
| **RgpA** | L215DRAFT_00128*<br>JCM13868DRAFT_01423[+] | 1.00E-06 | 26% | 308-502 | 1131 |
| **RgpB** | N/A | N/A | N/A | N/A | N/A |
| **Kgp** | L215DRAFT_00128*<br>JCM13868DRAFT_01423[+] | 3.00E-06 | 26% | 308-502 | 1131 |
| | L215DRAFT_00230*<br>JCM13868DRAFT_00677[+] | 2.00E-06 | 27% | 304-502 | 1131 |

*BLASTP hit in *P. uenonis* DSM23387 (IMG Genome ID 2528311143)

+BLASTP hit in *P. uenonis* DSM23387 (IMG Genome ID 2585427891)

[#]No BLASTP hits identified in *P. uenonis* 60-3

**Supplemental Table 4.** ***P. asaccharolytica*** **and** ***P. uenonis*** **proteins containing gingipain Pfams.**

| | | *P. asaccharolytica* | | *P. uenonis* | | |
|---|---|---|---|---|---|---|
| **Pfam ID+** | **Pfam Description** | CCUG 7834^ | PR426713P-I | CCUG 48615^ (IMG Genome ID 2528311143) | CCUG 48615 (IMG Genome ID 2585427891) | 60-3 |
| PF01364 | Peptidase family C25 | 2504824108# Poras_0230 | 650259179# | 2528766763# L215DRAFT_00971 | 2587120245# | 644453364# |
| PF03785 | Peptidase family C25, C-terminal ig-like domain | N/A | N/A | N/A | N/A | N/A |
| PF08126 | Propeptide_C25 | N/A | N/A | N/A | N/A | N/A |
| PF07675 | Cleaved Adhesin Domain | N/A | N/A | 2528765925# L215DRAFT_00128; 2528766027# L215DRAFT_00230 | 257119003# | 258711974# |
| PF10365 | Domain of unknown function (DUF2436) | N/A | N/A | N/A | N/A | N/A |
| PF18630* | Peptidase M60 C-terminal domain | N/A | N/A | N/A | N/A | N/A |

+Pfam IDs from *P. gingivalis* gingipains, RgpA, RgpB, Kgp

*Pfam only found in RgpB

#IMG/MER Gene IDs of Pfam hit

^Type strains used in this study

**Supplemental Table 5. Reciprocal BLASTP search of *P. asaccharolytica* and *P. uenonis* gingipain hits against *P. gingivalis*.**

| Strain | Query Gene ID/ Locus Tag | Top Blast Hits in *P. gingivalis* | % Identity | Query Coverage | E-value | Gene Name | Function |
|---|---|---|---|---|---|---|---|
| *P. asaccharolytica* CCUG 7834 | 250482108; Poras_0230 | PGN_0022 | 37% | 97% | 0 | PorU | Type IX secretion system peptidase |
| *P. uenonis* CCUG 48615 | 2528766763; L215DRAFT_00971 | PGN_0022 | 36% | 97% | 0 | PorU | Type IX secretion system peptidase |
| | 2528765925; L215DRAFT_00128 | PGN_1611 | 28.89% | 49% | 9.00E-41 | Leucine-Rich Repeat domain-containing protein | N/A |
| | | PGN_0852 | 29.11% | 34% | 9.00E-36 | Immunoreactive 47 kDa antigen | N/A |
| | | PGN_1733 | 27.10% | 17% | 2.00E-09 | Choice-of-anchor J domain-containing protein | N/A |
| | | PGN_1728 | 26.63% | 17% | 7.00E-06 | Kgp; Lys-gingipain | Gingipain |
| | | PGN_1155 | 24.14% | 12% | 1.00E-04 | Choice-of-anchor J domain-containing protein | N/A |
| | 2528766027; L215DRAFT_00230 | PGN_1733 | 25.37% | 33% | 1.00E-09 | Choice-of-anchor J domain-containing protein | N/A |
| | | PGN_1155 | 23.71% | 37% | 9.00E-07 | Choice-of-anchor J domain-containing protein | N/A |
| | | PGN_0335 | 28.92% | 13% | 2.00E-06 | PKD domain-containing protein | N/A |
| | | PGN_1728 | 24.77% | 37% | 1.00E-05 | Kgp; Lys-gingipain | Gingipain |

**Supplemental Table 6. Candidate collagenases in *P. uenonis* 60-3 and corresponding genes in *P. uenonis* CCUG 48615.**

| Accession ID (*P. uenonis* 60-3) | *P. uenonis* CCUG 48615* | Percent Identity | Protein Name | Protease Type | All Feature IDs in *P. uenonis* 60-3 | Differences in Collagenase Feature IDs^ |
|---|---|---|---|---|---|---|
| PORUE0001_1484 | L215DRAFT_00939 | 52% | Peptidase S1 domain-containing protein | Serine | IPR13783; IPR026444; TIGR0483 | N/A |
| PORUE0001_0731 | L215DRAFT_01109 | 72% | Trypsin | Serine | IPR13783; IPR026444; TIGR0483; SF49265 | IPR036116 (Fn-3 domain); SF49265; (Fn3 type III) |
| PORUE0001_1366 | L215DRAFT_01080 | 98% | Peptidase, S9A/B/C family, catalytic domain protein | Serine | SSF53474 | N/A |
| PORUE0001_0281 | L215DRAFT_00076 | 91% | Peptidase family M13 | Metallo | IPR024079 | N/A |
| PORUE0001_0568 | L215DRAFT_00076 | 93% | Peptidase family M13 | Metallo | IPR024079 | N/A |
| PORUE0001_0451 | L215DRAFT_00099 | 95% | Putative thiol protease/hemagglutinin | Cysteine | IPR026444; TIGR0483 | N/A |
| PORUE0001_1363 | L215DRAFT_01083 | 93% | Putative thiol protease/hemagglutinin | Cysteine | IPR026444; TIGR0483 | N/A |
| PORUE0001_1169 | L215DRAFT_01354 | 97% | Peptidase, S9A/B/C family, catalytic domain protein | Serine | IPR029058 | N/A |
| PORUE0001_1130 | L215DRAFT_01540 | 97% | Peptidase, U32 family | Unknown (U32) | PF01136; PF12392 | N/A |
| PORUE0001_0329 | L215DRAFT_00228 | 98.80% | Peptidase, U32 family | Unknown (U32) | PF01136 | N/A |

*Top BLASTP hit

^Present in *P. uenonis* 60-3 only
